# Genome dynamics in mosses: Extensive synteny coexists with a highly dynamic gene space

**DOI:** 10.1101/2022.05.17.492078

**Authors:** Alexander Kirbis, Nasim Rahmatpour, Shanshan Dong, Jin Yu, Nico van Gessel, Manuel Waller, Ralf Reski, Daniel Lang, Stefan A. Rensing, Eva M. Temsch, Jill L. Wegrzyn, Bernard Goffinet, Yang Liu, Péter Szövényi

## Abstract

**Background:** While genome evolutionary processes of seed plants are intensively investigated, very little is known about seed-free plants in this respect. Here, we use one of the largest groups of seed-free plants, the mosses, and newly generated chromosome-scale genome assemblies to investigate three poorly known aspects of genome dynamics and their underlying processes in seed-free plants: (i) genome size variation, (ii) genomic collinearity/synteny, and (iii) gene set differentiation.

**Results:** Comparative genomic analyses on the model moss *Physcomitrium (Physcomitrella) patens* and two genomes of *Funaria hygrometrica* reveal that, like in seed plants, genome size change (approx. 140 Mbp) is primarily due to transposable element expansion/contraction. Despite 60 million years of divergence, the genomes of *P. patens* and *F. hygrometrica* show remarkable chromosomal stability with the majority of homologous genes located in conserved collinear blocks. In addition, both genomes contain a relatively large set of lineage-specific genes with no detectible homologs in the other species’ genome, suggesting a highly dynamic gene space fueled by the process of *de novo* gene birth and loss rather than by gene family diversification/duplication.

**Conclusions:** These, combined with previous observations suggest that genome dynamics in mosses involves the coexistence of a collinear homologous and a highly dynamic species-specific gene sets. Besides its significance for understanding genome evolution, the presented chromosome-scale genome assemblies will provide a foundation for comparative genomic and functional studies in the Funariaceae, a family holding historical and contemporary model taxa in the evolutionary biology of mosses.

## BACKGROUND

The number of pseudomolecule-scale genome assemblies of seed plants has rapidly increased in the last 20 years revealing their conserved and divergent architectural features (1–5). In addition, comparative analyses of deep and shallowly divergent seed plant genomes provided detailed insights into genome evolution and dynamics both at longer and shorter timescales (6,7,16–20,8–15). By contrast, structure and dynamics of seed-free plant genomes are little understood (2,5,21). Comparison of the few available high-quality genomes suggests that overall genomic architectures of seed-free and seed plant genomes likely differ, which may be a consequence of their divergent genome dynamics (5,22–24).

Very little is known about the evolution of seed-free plant genomes, in particular over shorter timescales, mainly due to the lack of high-quality genome assemblies for groups of species with shallower genetic divergence. For instance, only five pseudomolecule-scale genome assemblies are available for the most intensively sequenced clade of seed-free plants, the mosses, but these are too deeply divergent to provide information on genome dynamics on a shorter timescale (23–26). Multiple aspects of seed-free plant genome dynamics remain unexplored. For example, the dynamics of genome expansion/contraction and structural variation are poorly understood and the contribution of transposable elements to these processes is debated (5,27–29). Also, little is known about the variation in gene content among species and whether it is shaped primarily by gene family diversification, genome duplication, or *de novo* gene gain/loss (30, 31). Preliminary data suggest that gene presence/absence variation may be common, nevertheless the contribution of gene family diversification may also be considerable. Furthermore, the genomic consequences of gene duplication, genome expansion/contraction and transposable element activity on genomic collinearity remain unexplored. To further our understanding about genome evolution and dynamics in seed-free plants, comparative genomic analyses of a group of relatively closely related species are needed.

Mosses compose the most species-rich lineage of bryophytes (mosses, liverworts and hornworts) and the group with the greatest number of pseudomolecule-scale genome assemblies among seed-free plants (23–26,32). Among mosses, the Funariaceae provide an appropriate model system to investigate genome evolution and dynamics in seed-free plants, as they hold the most widely used model organism of seed-free plants, the moss *Physcomitrium (Physcomitrella) patens*, for which a high-quality genome sequence and an ever-growing plethora of genomic resources are available (25,33–37). In addition, the Funariaceae exhibit broad diversity in morphology and development, habitat preferences, genome size, ploidy level and chromosome numbers spanning 90 million years of evolution (33,37–40). Finally, the family comprises shallow and more deeply divergent species whose evolutionary relationship has been extensively investigated in the last years (34,35,37,41). The considerable morphological and genomic diversity, the availability of the high-quality *P. patens* reference genome, and the intensively investigated phylogenetic backbone makes the Funariaceae a prime model to explore key questions of genome evolution in seed-free plants in general and in mosses in particular.

We present pseudomolecule-scale genome assemblies for two accessions of the moss *Funaria hygrometrica*, a further member of the Funariaceae, and compare these to the genomes of *P. patens* and other mosses. *F. hygrometrica* and *P. patens* diverged some 60 my ago (37) and differ considerably in their karyotype, genome size, gene content, and dispersal capability. In particular, (i) the sequenced accession of *P. patens* has 27 chromosomes (25, 42), while chromosome counts between 14-18 have been also reported for other isolates (42). By contrast, chromosomal races with 14, 26, 28, and 56 chromosomes have been reported for *F. hygrometrica* (25,39,43–46). Although chromosome numbers of some *F. hygrometrica* accessions and *P. patens* are similar (26/28 in *F. hygrometrica* and 27 in *P. patens*) their genome sizes considerably differ (*P. patens*: 511Mbp (c-value 0.53 pg); *F. hygrometrica*: 380 Mbp, 0.4 pg) (25,47–49). (ii) Preliminary data also suggest considerable gene content divergence between the two species, which may be related to their divergent habitat preferences and morphologies (30). Finally, (iii) genomic differences may also be driven by the different effective population size of the two species (50). Limited dispersal capability, high selfing and turnover rates of *P. patens* populations are expected to decrease species-wide effective population size, selection efficacy, and genome-wide genetic diversity (51–54). Therefore, *F. hygrometrica* and *P. patens* provide an appropriate species pair to investigate (i) the mechanism of genome size change, (ii) the contribution of gene gain/loss and duplication to gene content variation, (iii) and their overall effect on genomic collinearity in mosses, a diverse lineage of seed-free plants, in a simple haploid setting.

Our analyses consistently resolved 26 chromosomes in both accessions of *F. hygrometrica,* which can be easily derived by a chromosome break/fusion from the 27 chromosomes of *P. patens*. Like in seed plant genomes, the genome size difference between the two species (roughly 140 Mbp) was largely due to the expansion/contraction of transposable elements (TE) with no genomic hotspots. Despite similar gene numbers, the genomes of both species contained a large proportion (40%-30%) of species-specific genes that likely arose *de novo* while gene gain/loss through gene family expansion/contraction was less significant. Self- and between-species synteny revealed two whole-genome duplications, the older one is shared with various mosses whereas the more recent one is only shared with *P. patens*. Despite these dynamic changes, the *F. hygrometrica* and *P. patens* genomes retained remarkable synteny and collinearity following 60 million years of divergence. While synteny between chromosomes is maintained, inversion of hundreds of collinear blocks across the genome can be observed. Finally, genes and transposable elements showed rather uniform distribution across the chromosomes with no pericentromeric regions specifically enriched for TEs. This is in line with the hypothesis that large-scale genome structure of bryophytes and seed plants differ. Overall, our genome analyses suggest a genome structure in which rigid blocks of core genes coexist with a highly dynamic set of non-homologous genes leading to considerable gene content variation among genomes. Besides its contribution to understanding genome evolution in seed-free plants, our data will enable comparative analyses across the Funariaceae to investigate the genomic changes underlying the biological diversity at various scales.

## RESULTS

### *F. hygrometrica* accessions have 26 pseudomolecules

We assembled the genome of two *F. hygrometrica* accessions (one collected in Sankt Gallen, Switzerland [hereafter referred to as “Zurich”]; the other in Willimantic, Connecticut, USA [“UConn”]) using long-reads, Chicago and Hi-C libraries at the level of pseudomolecules (Supplementary_Table_1). Both assemblies were of high-quality resulting in 26 large scaffolds containing 99.10/96.11% (first 26 scaffolds Zurich accession=277486149 bp; first 26 scaffolds UConn accession=301785107 bp) of the approx. 300 Mbp (Full length of the assemblies: 280 Mbp Zurich, 314 Mbp UConn) genome with a minimum proportion of gaps for both accessions (Supplementary_Table_2). [[A browsable version of the genomes will be available upon acceptance]].

The assembled genomes were somewhat smaller than their estimated genome sizes using k-mer analysis or flow cytometry (Supplementary information). Whole-genome alignment and dot plot analysis of the genomes of the two accessions revealed highly collinear scaffolds (Figure 1 and Supplementary information), suggesting the absence of large-scale misassemblies in either of the assemblies and thus correspondence between the 26 largest scaffolds and the 26 putative chromosomes. These observations are in line with previous chromosome counts reported for *F. hygrometrica* (43,45,46,55). Contigs unanchored to the 26 pseudomolecules were short and contained few genes (Supplementary_Table_3). Despite being highly collinear, assembly length of the two accessions was different with the UConn accession genome being 34 Mbp longer, and the difference partially due to structural variation (Supplementary_Table_4 and Supplementary information). When aligning the assembled scaffolds of the two genomes with a similarity threshold of 50%, we found that 7% and 15% of the sequences were specific to the Zurich or UConn accession, respectively (Supplementary_Table_5). Nevertheless, accession-specific scaffolds were short and housed few if any genes (Supplementary_Table_3, for a more comprehensive comparison of the two accessions’ genomes see the Supplementary information).

**Figure 1.**
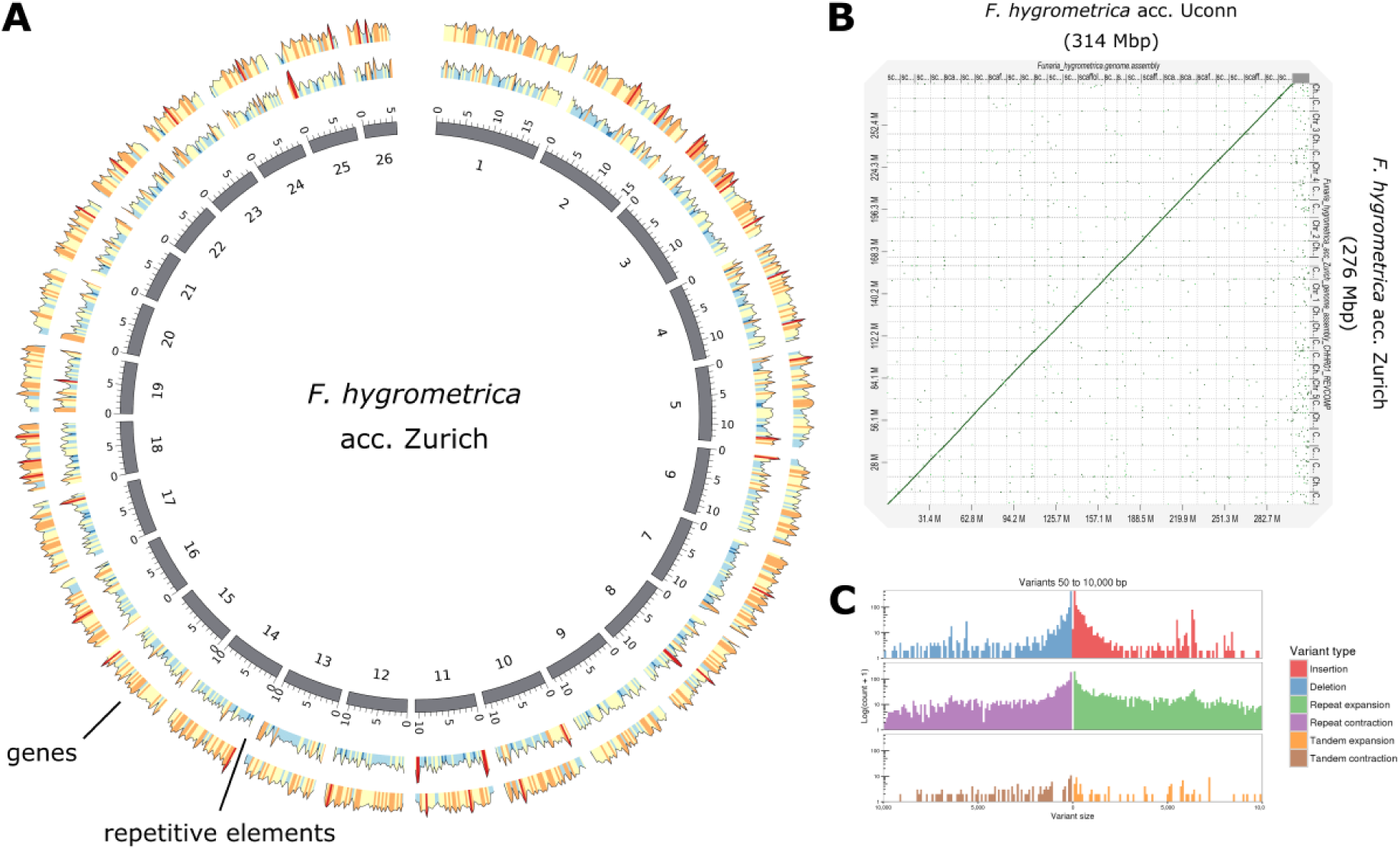
Organization and structural variation of the two *F. hygrometrica* genomes. (A) Circos representation of chromosome-scale pseudomolecules of the Zurich assembly. From outer to inner circle: Gene density in 250-kb windows along the putative chromosomes; Density of repetitive elements in 250-kb windows; representation of the 26 putative chromosomes (tick spacing is one Mbp). (B) Dot-plot of alignable regions between the UConn and Zurich assembly of the *F. hygrometrica* genome. The plot was generated with D-GENIES using a similarity threshold of 70% (145). (C) Length and frequency distribution of structural variants between the Zurich (reference) and UConn assembly. Variants were classified and plotted using assemblytics (148).

### Karyotypes of *F. hygrometrica* and *P. patens* are connected with a chromosome fusion/break

Both *F. hygrometrica* assemblies suggest the presence of 26 chromosomes in contrast to the 27 chromosomes reported for *P. patens*. Dot plots between the two genomes showed that all *P. patens* chromosomes have a distinct corresponding collinear chromosome in the *F. hygrometrica* genome (Figure 1 and 2, and Supplementary Information), except for Chr25 and Chr27 of *P. patens* (25) which both mapped to a single *F. hygrometrica* chromosome. This difference between the *P. patens* and *F. hygrometrica* assemblies was confirmed with both *F. hygrometrica* genomes and was not due to misjoins of the Hi-C scaffolding (see Supplementary Information). This implies that the chromosome number difference between *P. patens* and *F. hygrometrica* (27 and 26) can be either explained by a chromosome fusion (*P. patens* -> *F. hygrometrica*) or a chromosome break (*F. hygrometrica* -> *P. patens*) involving Chr27 and Chr25 of *P. patens* and Fh17 of *F. hygrometrica* (Figure 2).

**Figure 2.**
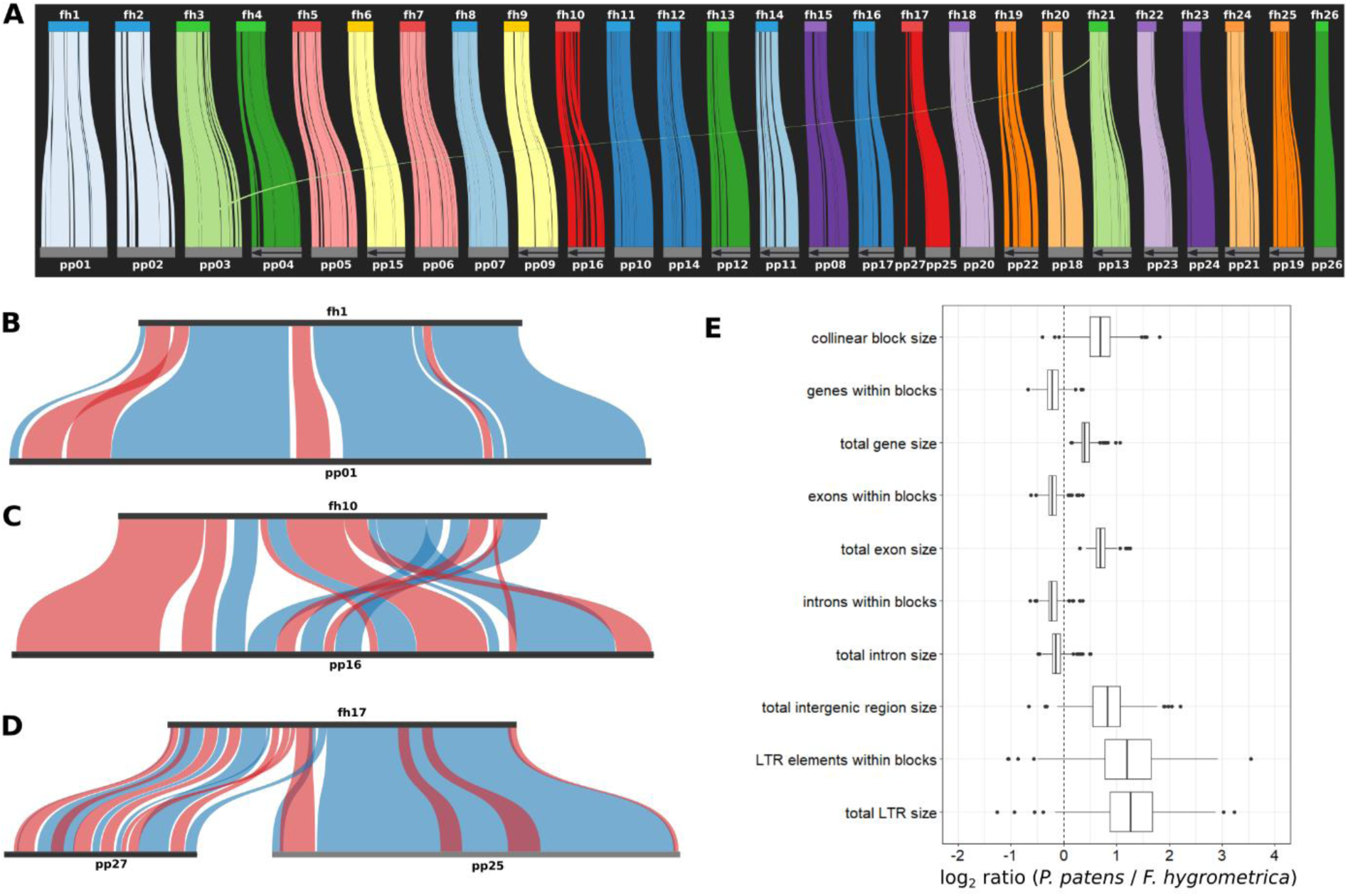
Intergenomic collinearity between the *P. patens* and *F. hygrometrica* acc. Zurich genome. (A) Collinearity between the *P. patens* and *F. hygrometrica* pseudomolecules. Collinear blocks of the two genomes are connected with colored lines and pseudomolecules are drawn to scale. (B-C) Collinearity between syntenic pseudomolecules of *F. hygrometrica* and *P. patens* (fh1 vs. pp1, and fh10 vs. pp16). Blue ribbons connect collinear blocks with the same directionality, while red ribbons depict blocks with inverted positions. (D) Collinearity and synteny between fh17, pp25 and pp27 representing the inferred chromosomal fusion/fission between chromosomes of *P. patens* and *F. hygrometrica*. (E) Comparison of genomic features in the corresponding collinear blocks of *P. patens* and *F. hygrometrica*. The box plots show the median and interquartile ranges, whiskers represent values up to 1.5 times the interquartile range, values outside this range are represented as individual data points.

### The *F. hygrometrica* genome is considerably smaller than the *P. patens* genome

Evidence from flow cytometry measurements (*P. patens*: 511 Mbp [c-value 0.53 pg]; *F. hygrometrica*: 380 Mbp, [c-value 0.4 pg], for further information see Supplementary information) and genome assembly results (assembly length: *P. patens*: 467 Mbp; *F. hygrometrica* Zurich accession: 280 Mbp; UConn accession: 314 Mbp) reveals that the *F. hygrometrica* genome is at least 130-150 Mbp smaller than the *P. patens* genome (Supplementary_Table_2, see also in Supplementary information) (25, 56). If genome size differences were due to random sequence gain/loss, we would expect that both species exhibit a similar proportion of species-specific sequence content. Alternatively, sequence gain/loss may have been asymmetric on the evolutionary branch connecting *P. patens* and *F. hygrometrica* with their common ancestor. We found that only 36% of the *F. hygrometrica* genome contained species-specific segments, whereas this fraction was almost twice as large (amounted to 62%) in the *P. patens* genome (Supplementary Information), suggesting accelerated sequence gain or sequence loss on the branch leading to *P. patens* or to *F. hygrometrica*, respectively. Genome size change has similarly affected each chromosome as was indicated by a positive correlation between chromosome lengths of *F. hygrometrica* and *P. patens* (Spearman’s Rho= 0.8611966, p≤ 1.884e-06). Assuming that the homologous portion of the genomes was inherited from the common ancestor, our finding suggests that the common ancestor of the two species and likely that of the Funariaceae was characterized by a smaller genome size than that of *P. patens*.

### Overall repeat content of the *F. hygrometrica* genome considerably differs from that of *P. patens*

The larger size of the *P. patens* genome may mainly be explained by the higher proportion of repeat elements. In line with this assumption, the nonalignable parts of the *F. hygrometrica* and *P. patens* genomes were enriched in their respective dominant LTR elements (see below), whereas the alignable segments were enriched in exonic and intronic regions (Figure 3A, Supplementary_Table_6). Nonalignable regions were also enriched in segments of the genomes containing no annotated features (regions outside of exons, introns, and repeat elements). The contribution of repeat expansion to the genome size increase was also supported by a significant positive correlation between the length and proportional TE content increase of homologous *P. patens* and *F. hygrometrica* chromosomes (Spearman’s Rho=0.517265, p-value = 0.00753). Altogether, this implies that the larger genome size in *P. patens* was primarily resulted from an increased representation of repeat elements and intergenic regions.

**Figure 3.**
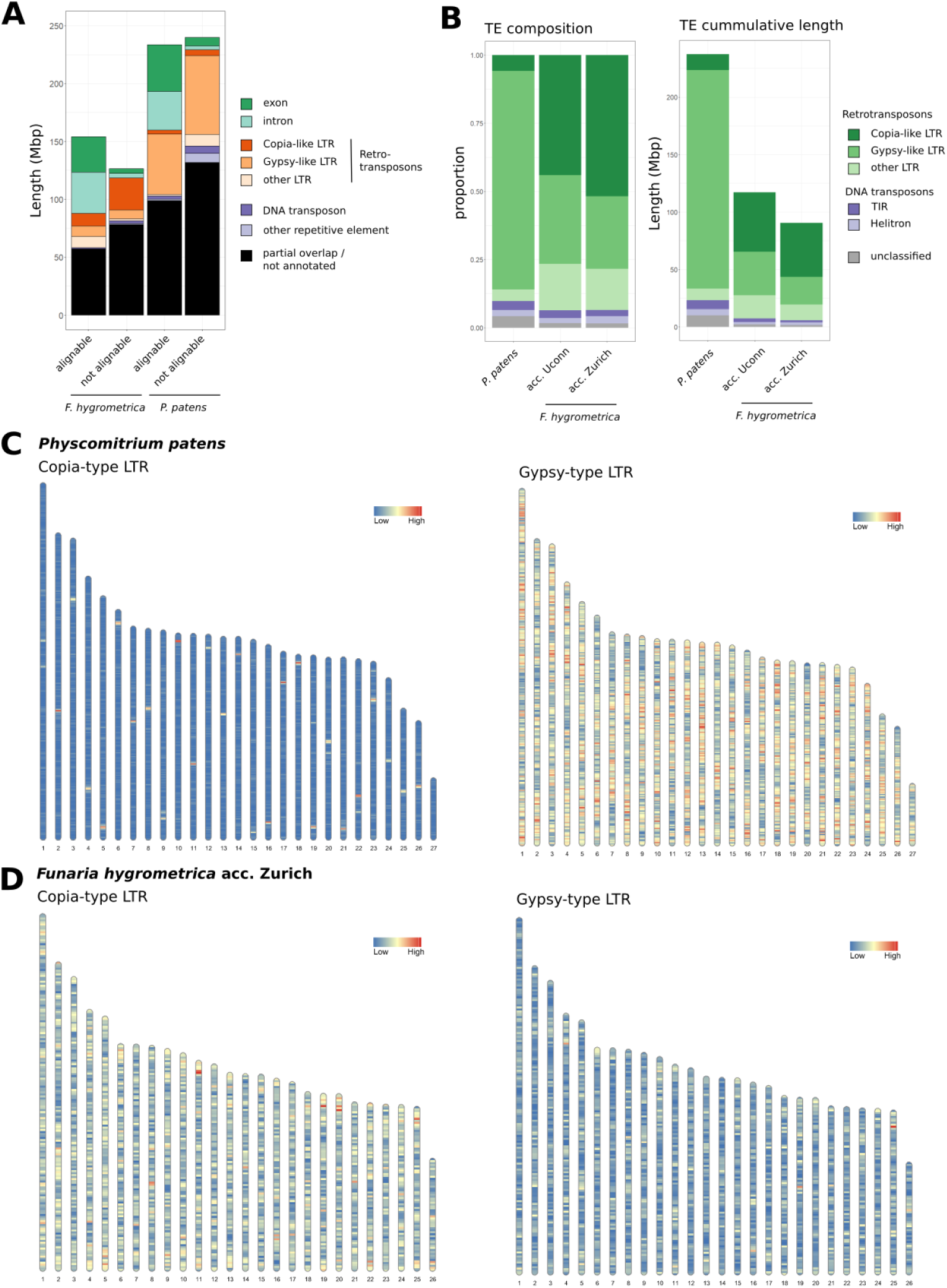
Transposable element annotation of Funariaceae genomes. (A) Genome features in alignable and not alignable regions between the *F. hygrometrica* acc. Zurich and *P. patens* genome. Only features fully overlapping with the respective genomic regions are shown as annotated, partially overlapping features are shown in black alongside regions without annotation. The whole-genome alignment was computed using the minimap2 algorithm with default parameters (146). (B) Transposable element annotation summary of the three studied genomes. The bar plot on the left shows the composition of transposable element families as a fraction based on their total sequence length in the genome. The plot on the right shows the absolute length the different families occupy in the respective genome. (C, D) Density of Long terminal repeat (LTR) retrotransposons on the chromosomal scaffolds of *P. patens* (C) and *F. hygrometrica* (D) in 100 kb windows.

A closer look at the repeat element content of the two genomes revealed that they differ both quantitatively and qualitatively. About a third of the *F. hygrometrica* genomes (32% and 37% of the Zurich and UConn accession’s genome, respectively) were predicted to be occupied by repeats (Figure 3B, Supplementary_Table_6). Compared to the *P. patens* genome, of which 51% are covered by repetitive elements, this amounts to a 13–18% difference in the fraction of repetitive elements between the *F. hygrometrica* and *P. patens* genomes. In other words, almost 80% of the genome size difference between *P. patens* and *F. hygrometrica* can be attributed to differences in repetitive element content alone. Furthermore, the class of LTR dominating the repeat content of the genome differed between the two species, namely Gypsy elements in *P. patens* (41% out of the total 51% repeat content) and Copia elements (16–17% [Copia] vs. 9–12% [Gypsy]) in *F. hygrometrica* (Figure 3B, Supplementary_Table_6).

The overall difference in repeat content and the differential abundance of Copia and Gypsy elements between the *F. hygrometrica* and *P. patens* genomes could have arisen by lineage-specific expansion of LTRs. Intriguingly, our reanalysis of the temporal activity of Copia and Gypsy element insertions in the *P. patens* and *F. hygrometrica* genomes revealed shared histories (Figure 4). While Copia elements exhibited rather continuous activity through time albeit with a recent and an older peak of activity in the genome of both species, and Gypsy elements having been active mainly in the recent past (Figure 4), the temporal dynamics of dominant LTR elements did not differ between the two genomes. Further, the absolute number of intact Gypsy elements was more than two-fold higher in the repeat rich and Gypsy-dominated *P. patens* than in *F. hygrometrica* (ca. 72000 [*P. patens*] vs. 16000/30000 [*F. hygrometrica* Zurich/UConn], see Table 1). The significantly greater number of all and intact Gypsy elements in *P. patens* and the similar proportion of intact elements in *P. patens* and *F. hygrometric*a suggest that the difference between the two genomes likely arose via a more massive activation of Gypsy elements in *P. patens*. By contrast, the activity of Copia and Gypsy LTRs was more balanced in *F. hygrometrica*, but overall at a lower level compared to *P. patens*.

**Figure 4.**
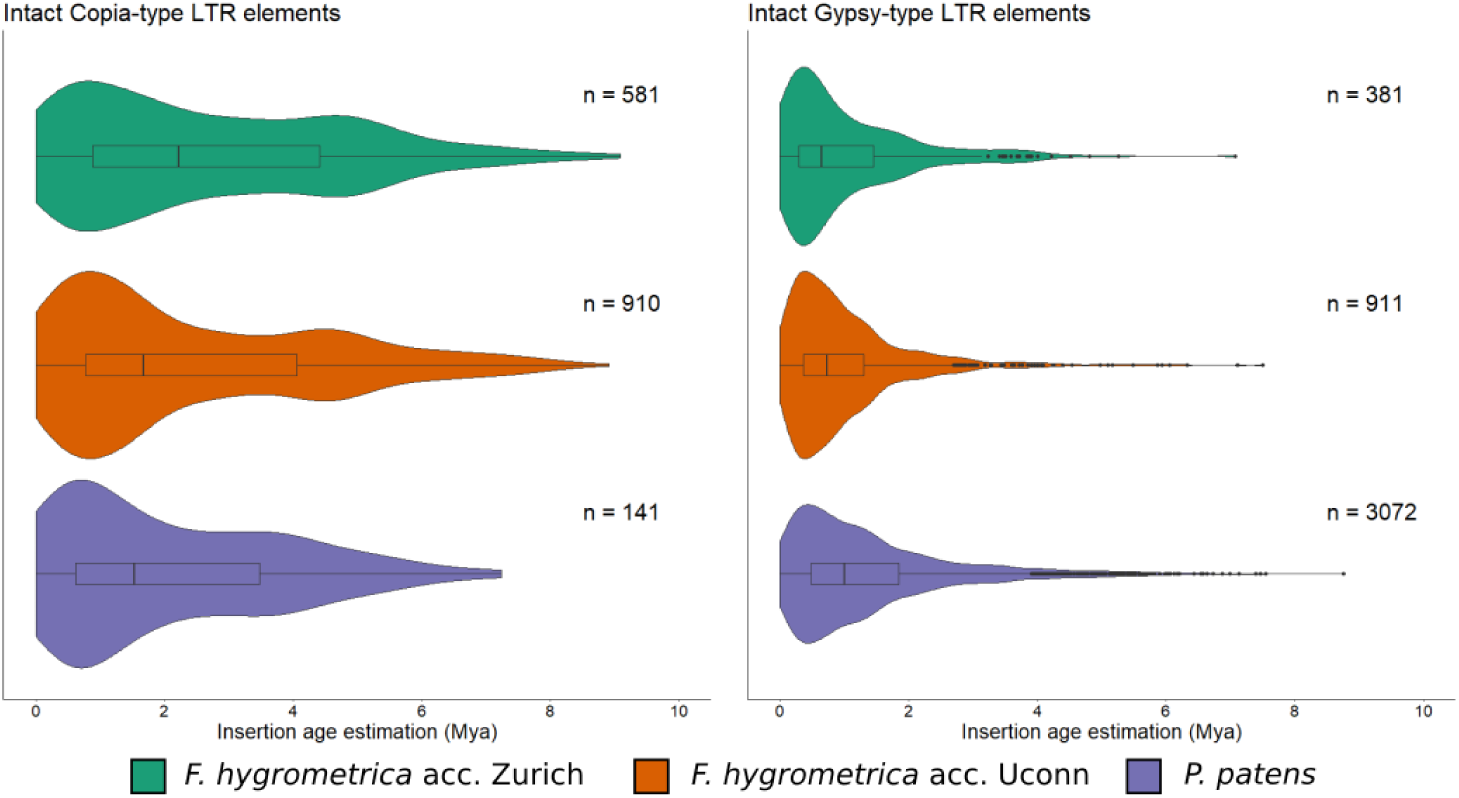
Insertion age distribution of intact full-length LTR elements in the Funariaceae genomes. Insertion time is estimated by calculating the sequence divergence between left and right terminal repeats and provided in millions of years (Mya). "n” refers to to the number of full-length elements included in the analysis..

**Table 1:** Proportional and absolute abundance of Copia- and Gypsy-like LTR elements in the *F. hygrometrica* and *P. patens* genomes as estimated by the EDTA pipeline (121).

|  | *F. hygrometrica*<br>acc. Zurich | *F. hygrometrica*<br>acc. Uconn | *P. patens* v3.3 |
| --- | --- | --- | --- |
| # Copia elements | 49,215 | 64,501 | 19,670 |
| # intact Copia elements | 27,990 (57 %) | 40,481 (63 %) | 13,451 (68 %) |
| # Gypsy elements | 24,801 | 69,216 | 187,270 |
| # intact Gypsy elements | 16,204 (65 %) | 30,697 (44 %) | 72,565 (39 %) |

### *F. hygrometrica* has higher gene density than *P. patens*

While genomes of the sequenced *F. hygrometrica* accessions were considerably smaller than those of *P. patens*, they harbored more genes. Our genome annotations resulted in 36,301 and 36,804 filtered gene models for the UConn and Zurich accessions of *F. hygrometrica*, respectively (Supplementary_Table_7), which is about 3,000 genes more than the predicted 32,926 genes in *P. patens* (25). Predicted gene sets of the *F. hygrometrica* accessions were of high-quality. Importantly, about 80% of the predicted gene models were supported by expression evidence (85% of Zurich and 74% of UConn gene models have RNAseq coverage higher than 80%) (Supplementary_Figure_1). Furthermore, BUSCO scores of both the annotated gene set and that of the genome sequence were among the top of currently published chromosome-scale genomes (including *P. patens*) (Supplementary_Table_8). Due to the smaller genome size and the larger gene set, gene density of the *F. hygrometrica* genome is nearly twice that of *P. patens* (13.08/12.20 genes/100 kbp [*F. hygrometrica* Zurich/UConn] vs 7.12 genes/100 kbp [*P. patens*]).

### Both the *F. hygrometrica* and *P. patens* genomes harbor a large proportion of species-specific genes

To compare homology of the gene set in the two species using a phylogenetic approach, we created orthogroups using proteomes of 38 plant species including 12 bryophytes, the two *F. hygrometrica* accessions, various vascular plants, green and streptophyte algae (Supplementary_Table_9). We recovered 52,231 orthogroups including 82.5% of the genes with only 0.5% of the orthogroups being species-specific (Supplementary_Table_10). The majority (40,966 or 78.43%) of these families contained bryophyte genes and more than half had at least one moss gene (30,510 or 58.41%). 44.51% (23, 248) of the gene families harbored genes for the three Funariaceae species (*F. hygrometrica, P. patens*, and *Physcomitrium pyriforme*) included in our analyses and 22,324 gene families contained at least one gene for our two focal species (*F. hygrometrica* and *P. patens*). Overall, 75.44% (24,839/32,926) of the *P. patens* and 90.01–90.41% (Zurich: 33,146/36,103; UConn: 33,274/36,804) of the *F. hygrometrica* gene models could be assigned to orthogroups (Supplementary_Table_10).

A significant proportion (41.11%, 15,026 genes) of the *F. hygrometrica* gene set occurs in lineage-specific gene families, compared to 30.5% (i.e., 10,044 genes) of the *P. patens* gene set. Therefore, shared gene families housed about 60% (58.89% 21,527/36,553) and 70% (69.50% 22,882/32,926) of the *F. hygrometrica* and *P. patens* gene sets, respectively (Supplementary_Table_10). Presence/absence polymorphism of genes was not an artifact of gene prediction. Virtually all predicted *F. hygrometrica* gene models had RNA-seq coverage (see Supplementary_Figure_1) and gene families with species-specific genes had genes predicted for both accessions (Supplementary_Table_10). Furthermore, only 4.92% (739 gene models) of the 15,026 *Funaria*-specific genes could be partially (50% coverage threshold) mapped to the *P. patens* genomic sequence of which 42.63% (315) produced truncated gene models with one or more frameshifts. *P. patens* holds a similar set of species-specific genes. Only 8.69% (873) of the 10,044 *P. patens*-specific genes had partial matches in the *F. hygrometrica* genome sequence of which 20.96% (183) were further interrupted by frameshifts. Thus, most of the lineage-specific genes cannot be detected in the alternate genome sequence, and therefore likely represent *de novo* gene gains/losses following the divergence of the two species. By contrast, a considerably smaller proportion of lineage-specific genes are likely due to gene degeneration/pseudogenization in one of the two genomes.

Although the proportion of genes unique to either of the two species was considerably high, over 60% of the gene set occurred in shared gene families (Supplementary_Table_10). Therefore, we also assessed how gene family expansions and contractions of shared gene families have contributed to the gene space of the two species (Supplementary_Table_11). Our Bayesian analysis indicated that only a few gene families have significantly expanded/contracted (posterior probability of expansion/contraction ≥ 0.8) on the branch leading to *F. hygrometrica* or *P. patens* from their most recent common ancestor. The proportion of shared gene families showing significant size change (posterior probability ≥ 0.8) was 0.91% (190/20,980) and 1.04% (219/20,980) for *P. patens* and *F. hygrometrica*, respectively. Furthermore, gene family evolution proceeded almost exclusively through expansions (i.e., 190 families) versus contractions (i.e., 0) in *P. patens*. By contrast, fewer families expanded (175 families) on the branch leading to *F. hygrometrica* and significantly more families were contracted (44 families). The gene set difference between *P. patens* and *F. hygrometrica* is therefore likely achieved primarily by *de novo* gain/loss of genes and not by gene family diversification.

### The *F. hygrometrica* and *P. patens* genomes retained high-level of collinearity

The *P. patens* genome appears to have significantly expanded via the activation of LTRs, which might have led to an extensive spatial reshuffling of the gene set. We therefore assessed the effect of LTR expansion on the collinearity of the two genomes. Despite considerable genome expansion and 60 million years of independent evolution, we found remarkable gene-level collinearity between the two genomes (Figure 2A and Supplementary_Table_12). More specifically, about half of the *F. hygrometrica* and *P. patens* gene set (49.52% or 17,977 genes and 55.08% or 18,137 genes in *F. hygrometrica* [Zurich accession] and *P. patens*, respectively) occurred in 845 collinear blocks containing at least five collinear genes. Knowing that about 70% and 60% of the *P. patens* and *F. hygrometrica* gene set has homologs in the alternate genome, this implies that almost all shared genes (80-90%) are found in collinear blocks. Despite the remarkable collinearity, inversions of collinear gene blocks were not uncommon between the two genomes. About half of the collinear blocks (51.95%, 439) were inverted. Nevertheless, inverted and noninverted collinear blocks had very similar genomic properties: they did not differ in their overall number of genes, number of collinear gene pairs and the genomic length of collinear segments in both genomes. Therefore, inverted regions did not serve as hotspots of genome evolution. Altogether, our observations indicate that despite new LTR insertions and 60 million years of divergence, collinearity was retained over most of the genome. Functional significance and the genomic/molecular mechanisms leading to this remarkable collinearity are unknown and must be explored.

### Smaller genome size of *F. hygrometrica* is mirrored by its shorter collinear regions compared to *P. patens*

Because the majority of the genome was covered by collinear gene blocks, we expected that genome size increase has led to expanded collinear blocks in *P. patens* compared to *F. hygrometrica* (Figure 2E and Supplementary_Table_12). In line with our expectation, the overall size of the genomic segments containing the collinear blocks was about twice as large in the *P. patens* compared to the *F. hygrometrica* genome (Fh_median_= 271,894 bp, interquartile range [IQR]= 166,567-455,659 bp; Pp_median_=444,906 bp, IQR= 260,162-779,820, Wilcoxon rank sum test W = 248333, p<2.2e-16). This difference was largely due to the increased size of intergenic regions in *P. patens* compared to *F. hygrometrica* (Fh_median_= 190,100 bp, IQR= 109400-328200; Pp_median_= 325,400 bp, IQR= 182000-617800), while genomic segments of the collinear blocks contained somewhat fewer genes in *P. patens* than in *F. hygrometrica* (Pp_median_=34.00, IQR=22.00-56.00; Fh_median_= 39.00, IQR=25-64; W = 110432, p<2.2e-16). This is in line with our previous assertion that genome expansion was primarily achieved by repeat expansion leading to an overall decreased gene density in collinear blocks in *P. patens* versus in *F. hygrometrica* (Figure 2E).

### The most recent whole genome duplication is shared by *F. hygrometrica* and *P. patens*

Previous analyses suggested that the ancestor of mosses may have had seven chromosomes, which then underwent two whole genome duplications (WGD) in *P. patens* (23–25). Signatures of the older whole-genome duplication dated to about 200 mya were shown to be shared by *Ceratodon purpureus* (23, 57) and *Syntrichia caninervis* (24), whereas the more recent one likely predated the origin of the Funariaceae (25,30,38,41,58). We found abundant collinearity and synteny between the *P. patens* and *F. hygrometrica* chromosomes (Figure 2A-D), and our Ks analysis resulted in very similar Ks distributions in *F. hygrometrica* and *P. patens* (Figure 5). Furthermore, both species’ Ks distribution showed two major peaks at Ks=∼0.8 and Ks=∼1.2 representing two potential WGDs. Self-synteny maps of both genomes were also very similar, further confirming the presence of two shared WGDs (Figure 5). Therefore, our Ks and self-synteny-based analyses suggest that both the old and the more recent whole-genome duplications are shared by the two species (Figure 5). This confirms that both WGDs preceded the split of *F. hygrometrica* and *P. patens* and that the more recent WGD represents a Funariaceae-wide and potentially Funariaceae-specific duplication event.

**Figure 5.**
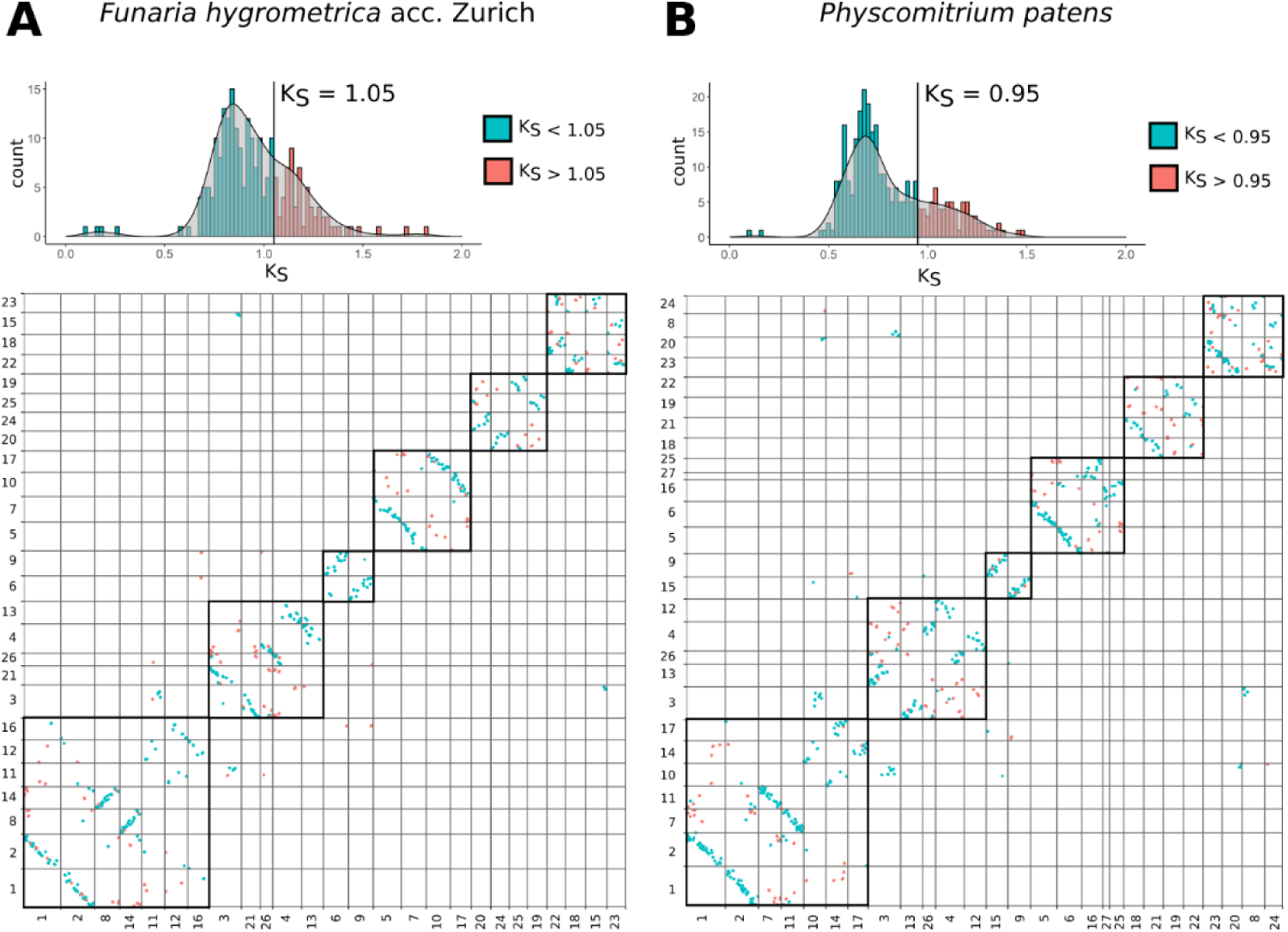
Dot plot of self synteny among pseudomolecules of the *F. hygrometrica* Zurich accession and *P. patens*. Pseudomolecule blocks corresponding to the putative ancestral chromosomes are framed. Histograms above the dot-plots show the distribution of average Ks values per collinear block. Histograms are colored according to the two whole-genome duplication events. Pseudomolecules in the dot plots are ordered according to intergenomic synteny between *F. hygrometrica* and *P. patens*.

### Overall chromosome structure of *F. hygrometrica* resembles that of *P. patens* and other bryophytes

The overall chromosome structure of the *F. hygrometrica* genomes is very similar to that of the other published bryophyte genomes. Pericentromeric regions enriched for TEs could not be identified and gene and repeat features were rather uniformly distributed along the chromosomes (Figure 1A, Figure 3C-D). Although pericentromeric regions enriched in TEs are not present in the *P. patens* genome, RLC Copia elements show a peak at the centromeric regions (25). In the *F. hygrometrica* genome, we could not identify a single Copia or Gypsy subfamily that showed a clear and single peak in each pseudomolecule (Figure 1 and Supplementary Information). Therefore, we conclude that association of RLC elements with the putative centromeres is specific to *P. patens* and does not occur in *F. hygrometrica*.

## DISCUSSION

Comparison of sequenced seed-free and seed plant genomes suggests that their overall genome structure may differ in multiple aspects (5) yet given the paucity of information on the genome evolution and dynamics of seed-free plants, the ultimate causes of their differential genome structure is poorly understood (21). Here, we present two new pseudomolecule-scale genome assemblies for two accessions of the moss *F. hygrometrica*, a relative of the most often employed seed-free model *Physcomitrium* (*Physcomitrella*) *patens* to investigate genome evolution and dynamics in the most diverse group of seed-free plants, the mosses, in a simple haploid setting. More specifically, by conducting comparative analyses of the *F. hygrometrica* and *P. patens* genomes, we focused on three major aspects of genome evolution: (a) the genomic processes underlying genome size change, (b) the degree and evolution of genomic collinearity, and (c) the processes contributing to gene set differentiation. We discuss these results and their genome evolutionary implications in the following paragraphs.

Genome size is an important characteristic of organisms significantly affecting their short- and long-term evolution (59–63). In many seed plants, genome size variation is caused by the expansion/contraction of non-coding DNA, especially TE-elements (64–66). Nevertheless, exceptions to this rule are known and the number and type of TE-families contributing to genome size change varies among lineages of seed plants (67). By contrast, direct genomic evidence for the predominant effect of TEs in the genome size variation of bryophytes such as mosses is lacking. We provide evidence that the 150 Mbp larger genome size of the *P. patens* (versus the *F. hygrometrica*) genome is primarily due to its increased TE content, suggesting that genome size variation in the Funariaceae, and hence perhaps in mosses and bryophytes in general, is driven, like in seed plants, by expansion/contraction in TE-elements. While very little indirect evidence is available for other mosses to support this hypothesis, cytological data on liverworts suggests the presence and expansion of large heterochromatic regions in taxa with larger genomes (68, 69). Further studies will be needed to test the generality of this observation in mosses, bryophytes and other seed-free plants.

While the contribution of TEs to genome size variation is evident, the driving factors of genome size variation remain unclear. In flowering plants effective population size is likely one of the most important factors affecting genome size evolution (70). Both *P. patens* and *F. hygrometrica* are monoicous moss species capable of simultaneously producing genetically identical motile sperm cells and sessile egg cells on the same haploid plant (gametophyte) (33). Therefore, fertilization often occurs through the union of the genetically identical gametes (intragametophytic selfing) resulting in a fully homozygous diploid sporophyte (71–73). In such sporophytes, spore formation via meiosis resembles clonal reproduction because spore progeny is expected to be genetically identical. Intragametophytic selfing is expected to severely decrease effective population size leading to an overall reduction of selection efficacy (53, 54). Intragametophytic selfing is thought to be more frequent in *P. patens* with a sunken spore capsule, lacking active opening mechanism, containing large and heavy spores preventing efficient spore dispersal via air currents. This is assumed to cause a greater decrease in effective population size and less effective purging of TE-elements in *P. patens* compared to *F. hygrometrica* (50,71,72). Therefore, the greater abundance of TEs is likely caused by the severely reduced effective population size of *P. patens* leading to a less effective control over the activity of TEs.

Previous comparative analyses among moss genomes that diverged over 170-200 million years ago revealed detectable synteny among chromosomes (so-called “ancestral elements”), representing conserved blocks inherited from the common ancestor of all mosses (23, 24). Consequently, synteny and collinearity should be even more pronounced between more recently diverged species, a hypothesis virtually unexplored in seed-free plants (26). Our analysis recovered unexpectedly strong collinearity between the *F. hygrometrica* and *P. patens* genomes, despite 60 million years of divergence (37). More specifically, almost all genes (roughly 80%) with homologs in the alternate genome were in collinear blocks. Furthermore, collinear blocks were also syntenic, showing virtually no movement of collinear blocks among chromosomes. This level of collinearity and synteny represents a relatively rigid genome structure that is exceptional compared to the data available for highly collinear grass and angiosperm genomes with similar depth of divergence (74–80) and is in line with findings by (26). While gene order, gene content, and chromosomal position of collinear blocks are highly conserved, their orientation appears to be dynamic. Indeed half of the collinear blocks are inverted between *F. hygrometrica* and *P. patens*. Therefore, structural dynamics of chromosomes in these two moss genomes is primarily driven by frequent inversion of highly stable collinear blocks within chromosomes. The evolutionary forces maintaining this extensive collinearity despite WGD and TE-expansion/contraction are currently unknown.

Despite extensive collinearity of homologous gene copies, we confirmed previous hypotheses that both the *F. hygrometrica* and the *P. patens* genome contain a large proportion of species-specific (lineage-specific) genes (30, 81). Our analyses also show that these species-specific genes have no detectable homologs in the other species’ genome and therefore likely arose *de novo* or emerged as specific following the loss through deletion or excision of the homolog in the other species’ genome. Finally, we also clarify that the observed gene content difference is not an artifact of the annotation process. Together, these observations suggest that gene birth/death has considerably contributed to the genome evolution of the Funarioid mosses. While the presence of lineage-specific genes is not surprising, their relative contribution to the gene space of each species is exceptionally large, reaching 20–30%. In comparison, gene space difference between highly contiguous angiosperm genomes with similar depth of divergence is less pronounced (17,80,82–89). For instance, the proportion of species-specific genes usually remains below 3–10% in most studies. This suggests that *de novo* gene birth/death may be more prevalent in mosses than in vascular plant genomes. While the mechanisms of gene birth/death are unclear, they may be linked to the activity of TEs, the shifting of reading frames, and/or pervasive transcription of various genomic regions (90–93).

Our observations allow us to provide a putative graphical model describing genome evolution and dynamics between *F. hygrometrica* and *P. patens* and potentially the Funarioid mosses. This differs in two main aspects from the genome dynamics observed in vascular plants: (i) stronger conservation of synteny and collinearity, and (ii) elevated rate of gene birth/death. More specifically, our observations suggest that rigid collinear homologous gene segments coexist with a highly dynamic non-homologous gene set with potential functional significance. While collinear homologous gene segments are kept together and their chromosomal order is mostly preserved, their orientation can change frequently. By contrast, lineage-specific genes are randomly dispersed across the genome and arise in a punctuated manner. The mechanisms and/or constraints driving this genome dynamic is unclear but may be related to the haploidy of the moss genome directly exposing mutations to natural selection. This is expected to increase the efficacy of purifying selection potentially leading to extended synteny and collinearity (54, 73). It is also possible that high efficiency of homologous recombination facilitates homology-mediated repair, which may increase genomic synteny and collinearity (25,33,94). Finally, elevated collinearity and synteny could also be linked to the relatively small and less variable genome size of mosses compared to flowering plants (49, 59). Nevertheless, the ultimate factors governing genome dynamics in mosses are unknown and need to be further explored. Similarly, the contribution of genomic changes to non-adaptive/adaptive variation within the Funarioid mosses are not known and need to be investigated.

All bryophyte genomes studied so far are characterized by an unusual chromosome structure with repeat and gene features relatively evenly spread along the chromosomes. While specific TEs may form a narrow peak in the middle of the centromeres, bryophyte chromosomes lack a broad TE enriched pericentromeric region typical for flowering plants (*Marchantia polymorpha* (95), *Ceratodon purpureus* (96), *Syntrichia caninervis* (24), *Anthoceros agrestis* (27)). This is in stark contrast to the usual chromosome structure of flowering plant genomes where gene density is highest in the middle of the chromosome arms whereas repeats are more dominant in the pericentromeric regions. It was proposed that this unique large-scale chromosome structure may be a feature of most bryophyte and seed-free plant genomes (5). Nevertheless, early cytological studies described the occurrence of chromocenters in *F. hygrometrica,* which can also be observed in the liverwort *M. polymorpha* but not in *P. patens* suggesting that overall chromosome structure of *F. hygrometrica* may differ from that of *P. patens* (25,97,98). Our study corroborates the hypothesis that moss and potentially most bryophyte genomes show the above-described unique genome structure. In addition, it also provides further evidence that not all moss and bryophyte genomes accumulate specific TE-elements in their putative centromeric regions. We note that similarly to *F. hygrometrica*, no dominant TE peak was found on the *S. caninervis* (24) and *A. agrestis* pseudomolecules (27) but specific TE families were colocalized with the putative centromeres of *C. purpureus, M. polymorpha*, and *P. patens* (23,25,97,99). It is currently unclear what processes shape the accumulation of a specific TE class at the putative centromeres in some (*P. patens*, *M. polymorpha*, and *C. purpureus*) but not in other bryophyte species. Detailed comparative analysis of the putative pericentromeric/centromeric regions of *F. hygrometrica* and *P. patens* may provide further insights into this question.

Based on the *P. patens* chromosome-scale genome assembly, an earlier study reconstructed the possible trajectory for karyotype evolution in mosses (25). According to this scenario, the first whole genome duplication (WGD) of ancestral chromosomes resulted in 14 chromosomes, which was followed by one chromosome loss and the fusion of another two chromosomes for a final karyotype of 12 chromosomes. This hypothesis is also fully supported by our collinearity analysis between the *F. hygrometrica* and *P. patens* genomes (Figure 6). The second WGD led to 24 chromosomes of which the 27 chromosomes of *P. patens* were derived by five breaks and two chromosome fusions. This scenario may also underlie the history of the karyotype of 26 chromosomes in *F. hygrometrica*, except the trajectory involving the origin of chr25 and chr27 in *P. patens*. This is because both *F. hygrometrica* assemblies support the presence of 26 and not 27 chromosomes as reported for *P. patens*.

**Figure 6.**
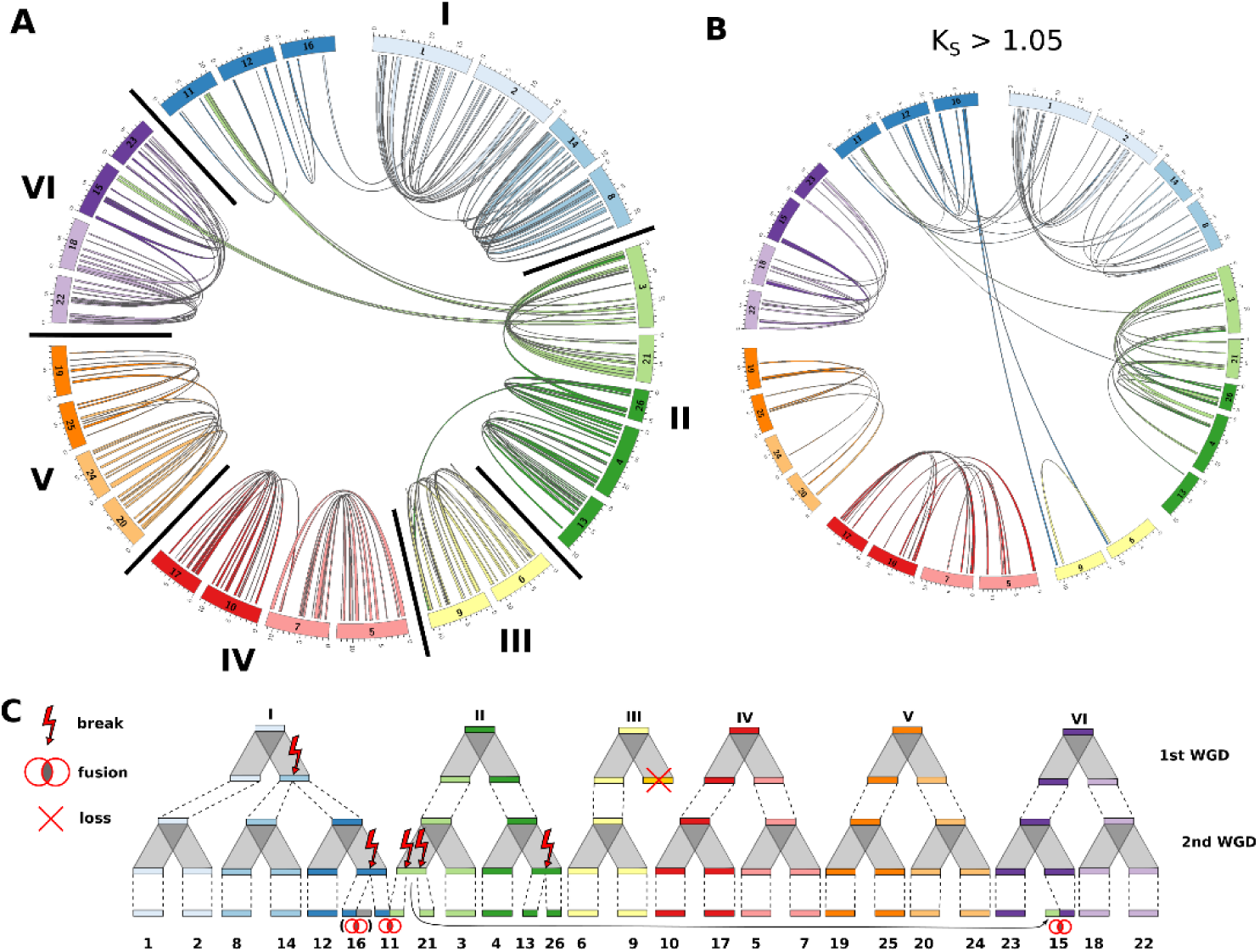
Intragenomic collinearity and karyotype evolution model of the *F. hygrometrica* genome. (A) Circular visualization of the 26 pseudomolecules of the *F. hygrometrica* genome (Zurich accession). Blocks of collinear genes with a mean synonymous substitution rate (Ks) ≤ 1.05 (corresponding to the most recent whole-genome duplication [WGD]) are connected by colored ribbons. Pseudomolecules are arranged to reflect their putative evolutionary relationship. Pseudomolecules potentially originating from the same ancestral chromosome are grouped together and labelled with I – VI. (B) Visualization of collinear blocks with a mean synonymous substitution rate (Ks) > 1.05 (corresponding to the older WGD). (C) Hypothetical model of karyotype evolution in the *F. hygrometrica* lineage. Six ancestral chromosomes underwent two whole genome duplication events accompanied by one chromosome loss, five chromosome breaks, and three chromosome fusions, resulting in 26 recent chromosomes.

An alternative and equally likely scenario could be based on an ancestral chromosome number of six Both six and seven ancestral chromosomes are in line with the chromosome number counts available for mosses (55, 100). This alternative scenario would involve one loss and one chromosome break after the first WGD and four breaks and three fusions after the second WGD (Figure 6). Nevertheless, available data are insufficient to distinguish between these two alternative scenarios.

Finally, the two pseudomolecule-scale *F. hygrometrica* assemblies also raise some questions concerning the accuracy of the current *P. patens* genome assembly and its actual chromosome number. Our study clearly implies that the *F. hygrometrica* accessions have 26 pseudomolecules, which can be easily derived from the 27 pseudomolecules of the *P. patens* genome by fusing two pseudomolecules (Chr25 and Chr27). Nevertheless, previous cytological studies have reported *P. patens* accessions with 27 as well as 26 chromosomes (45,46,55). Therefore, the possibility that the sequenced accession of *P. patens* had 26 chromosomes and that Chr27 and Chr25 represent falsely split segments of a single *P. patens* chromosome cannot be ruled out. In line with this hypothesis, putative centromeres of all *P. patens* pseudomolecules except Chr27 show a unique accumulation of RLC elements (25). Unfortunately, like Chr25 and Chr27 none of the *P. patens* chromosomes bear characteristic telomeric repeats, which could be used to trace their potential fusion. Furthermore, neither Hi-C library nor extensive long-read data are available for *P. patens* to resolve this issue and clarify the karyotypic changes between *F. hygrometrica* and *P. patens*.

## CONCLUSIONS

In comparison to the rapidly growing understanding of genome evolution and dynamics in flowering plants, very little is known about patterns and processes pertaining to changes in the genomes of seed-free plants (2, 5). Here, we sequenced and analyzed genomes of the moss family Funariaceae to investigate their evolution in the most speciose groups of seed-free plants, the mosses. Our analyses and the integration of previous observations suggest that moss genomes show more extensive synteny/collinearity and greater rate of gene birth/death than those of flowering plants. Therefore, our results provide further support to the hypothesis that genome dynamics of moss, bryophyte, and potentially seed-free plants differ from those of seed plants (5, 26). Our study provides a solid basis for a more extensive exploration of genome dynamics within the Funariaceae, to test for the generality of our observations. Moreover, availability of a high-quality genome sequence for two species representing end points of the morphological and ecological diversity within the Funariaceae will open the way for detailed investigations on the genetic basis of phenotypic diversity within the family (30,33–35,41,81).

## METHODS

### DNA sequencing

For both the Zurich and UConn accessions (Supplementary_Table_1) high molecular weight DNA was extracted using a modified CTAB protocol (101). The Genome of the Zurich accession was sequenced with Illumina and PacBio technology. Illumina libraries were generated with insert sizes of 250 bp, 350 bp, 2 kbp, and 5kbp and sequenced on Hiseq 2000, Hiseq 2500, and Hiseq 4000 systems (paired-end, 150 bp read length). PacBio data was generated on the RS II platform using C1 chemistry (3 cells) and P6-C4 chemistry (10 cells). Illumina sequencing yielded over 62 Gbp raw sequencing data in total, while PacBio sequencing resulted in 13 Gbp of sequence data.

For the UConn accession we generated two Illumina libraries with an insert size of 400 bp and sequenced them using the HiSeq Xten platform (paired-end, 150 bp read length). Using the very same DNA, we also prepared a single DNA library for Oxford Nanopore sequencing using the ligation kit and sequenced it on the Nanopore X5 platform. Illumina sequencing resulted in a total of 62 Gbp raw data. Nanopore sequencing resulted in about 1.8 million reads longer than 10,000 bp after clean-up.

To improve continuity of the genome assemblies we created Hi-C libraries for the Zurich accession (using the Dovetail Hi-C kit for genome assembly) and for the UConn accession using the protocol described in (102). Furthermore, a Chicago library was also prepared by Dovetail Genomics for the Zurich accession. We sequenced the Hi-C and CHiCAGO libraries using Illumina Hiseq 4500 and Novaseq machines in paired-end mode (150 bp read length). Details of the DNA sequencing data used for the genome assembly can be found in Supplementary_Table_1.

### Genome assembly

For the Zurich accession the initial assembly was generated with the Canu assembler v1.5 (103) combining all available PacBio data. Afterwards, we employed HiRise (104) together with the CHiCAGO sequencing data to scaffold the original reads into larger scaffolds and improve assembly contiguity.

The resulting assembly was further consolidated by a second HiRise run employing the Hi-C data. For manual curation, Hi-C sequencing data was aligned to the assembly using the juicer pipeline (105) and files for visual inspection of the Hi-C contact map were created with scripts supplied with the 3D-DNA software package (106). Visual review with the Juicebox software (107) revealed no obvious misassemblies, but we identified a misjoin in the largest pseudo-molecule. We manually corrected the misjoin and the genome assembly was updated to accommodate for the introduced scaffold split (see Supplementary Information).

For the UConn accession Nanopore raw reads were first corrected by Canu v1.9. The corrected reads were then assembled into contigs by NextDenovo v2.3.0 (https://github.com/Nextomics/NextDenovo) with default parameters. After assembly polishing (see below), Hi-C raw reads were processed by Juicebox v1.6 to extract valid reads which contain Hi-C contact information. The 3D-DNA pipeline was then used to cluster, orient, and order the contigs, generating chromosome-scale scaffolds. We also used Juicebox to manually adjust the scaffolding according to the contact map. After manual curation in Juicebox, the post-process module of 3D-DNA pipeline was used to generate the corrected chromosome-level scaffolds (see Supplementary Information).

### Polishing

The assembly of the Zurich accession was first polished with the quiver tool, which is included in the PacBio SMRT Analysis software package v2.3.0.140936, using all PacBio reads obtained with the P6-C4 chemistry. We used the default thresholds to remove very low coverage scaffolds from the assembly. This polishing step also corrected base calls, filled in Ns, and corrected repeat regions. After that we mapped Illumina reads to the quiver-polished assembly using BWA mem (108) to correct indels and SNPs in the non-repetitive parts of the genome assembly using Pilon v1.23 (109) in three rounds. Final polishing was done with PBSuite v.15.8.24 (110) to fill up some of the remaining gaps of the assembly using all available PacBio reads.

To fix SNPs, indels and SVs originated from sequencing errors, the UConn genomès assembly was first corrected using all Oxford Nanopore reads and the algorithm provided in racon v1.4.10 (111). We further corrected the racon polished contigs using all Illumina reads and Pilon v1.23. We repeated the successive polishing with racon and pilon three times. We used both tools with default parameters.

### Contamination detection and filtering

Initial assessment of the Hi-C contact maps of the Zurich accession suggested that some scaffolds of the assembly had very low coverage of Hi-C data and therefore represented potential contaminations. We made similar observation using the UConn accession’s genome assembly. To identify contaminant scaffolds, we used BlobTools v1.1.1 (112). More specifically, we used all Illumina reads and the full NCBI nucleotide collection (nt) and the uniprot database to assess sequencing depth along the genomic scaffolds and to assign them to broad taxonomic categories, respectively. After that, we removed all scaffolds from the assembly that were assigned to bacterial or other non-eukaryotic taxonomic classes.

### Repetitive element identification and annotation: RepeatModeler2

We used the automated approach implemented in the RepeatModeler2 package v2.0 (113) to generate a *de novo* annotation of repetitive elements in the genomes. RepeatModeler2 combines results from the repetitive DNA sequence discovery algorithms RepeatScout v1.0.6 (114) and RECON v1.08 (115) to generate a non-redundant set of TE families. Additionally, the RepeatModeler2 pipeline employs LTRharvest (116), which is included in the GenomeTools library v1.6.1 (117), and LTR_retriever v2.9.0 (118) for discovery of LTR elements based on structural parameters. To get a comprehensive annotation of repetitive elements, we first identified TE families present in the genome using RepeatModeler2 in conjunction with version 3.1 of the Dfam database of repetitive DNA families (119). The resulting library of TE families was then used for annotation of repetitive elements in the genome sequence using RepeatMasker v4.1.0 (120) (Supplementary_Table_13-14A and B).

### Transposable element annotation

We used the Extensive *de-novo* TE Annotator (EDTA) pipeline v1.9.6 (121) to get a comprehensive TE annotation for both *F. hygrometrica* genome assemblies. The pipeline combines output from several TE prediction and classification programs and applies a series of filtering steps to construct a comprehensive library of transposable elements present in the assembled genome (121). The contamination-filtered pseudomolecule-scale assemblies were used as an input to the pipeline together with coding sequences of genes annotated by BRAKER v2.1.0 (122) and genBlastG v1.39 (123) as described in the “Gene prediction” paragraphs. To avoid introducing a bias when comparing TE composition and distribution between the *F. hygrometrica* accessions and *P. patens* caused by differing annotation pipelines, we retrieved the most recent *P. patens* genome assembly and gene annotation v.3.3 (25, 47) and re-annotated transposable elements using the EDTA pipeline as described above (Supplementary_Table_15-17).

### Phylogenetic analysis of LTR Copia and Gypsy super-families

For further sub-classification of annotated LTR elements of the Copia and Gypsy super-family, we retrieved alignments of reverse transcriptase (RT) domain of several known sub-families from the *Gypsy* Database (124) and built a Hidden Markov Model (HMM) for the protein domain employing the hmmbuild function of the HMMER software package v3.3 (125). We then translated nucleotide sequences of Copia and Gypsy elements annotated in the *F. hygrometrica* genome to their respective peptide sequences in all six frames and scanned them for the presence of an RT domain in combination with the previously built HMM using the hmmsearch utility (126) of the HMMER software package v3.3 (125). We retained significant hits (E-value threshold: 1e-5) covering at least 80% of the protein domain. We discarded all LTR elements which had multiple valid hits in different reading frames. The remaining RT domains were aligned to each other with MUSCLE v3.8.31 (Edgar 2004) using the consensus sequence of the RT domain of the Bel-Pao superfamily, retrieved from GypsyDB, as outgroup. The phylogenetic tree of the RT domains was inferred using the Neighbor-Joining method (128) in MEGA X v10.2.5 (129) applying 1000 bootstrap replicates (130) (Supplementary_Information).

### LTR insertion time estimation

The sequences of 5‘ and 3’ terminal repeats are supposed to be identical to each other when LTR retrotransposons are newly inserted into the genome (131). Therefore, the degree of sequence conservation between left and right terminal repeats can be used as a proxy for insertion age of individual LTR elements. To assess the recent history of LTR retroelement insertions in *F. hygrometrica* and *P. patens*, we extracted 5’ and 3’ terminal repeat sequences for each LTR element classified as full-length and intact by the EDTA pipeline (121) and aligned them using MUSCLE v3.8.31 with default parameters (127). We then calculated the Kimura 2 parameter distance (132) for each aligned pair using a custom python script and modules from the Biopython library (133). The divergence time between LTR pairs was estimated by dividing the distance parameter by two times the synonymous substitution rate. We used a substitution rate of 9.4e-9 synonymous substitutions per synonymous site per year for both genomes, which was established for *P. patens* elsewhere (134). We plotted LTR insertion age distributions using the R package ggplot2 v3.3.3 (135).

### Genome annotation

#### Transcriptome assembly

To aid gene prediction we generated RNA-seq data covering three developmental stages of the gametophyte and four developmental stages of the sporophyte generations in three replicates (six developmental stages in total) for the Zurich accession. Gametophyte and sporophyte RNA-seq data was also obtained for sporophyte and gametophyte tissues of the UConn accession (Supplementary_table_1). RNAseq data was first trimmed for low quality bases and adapter sequences using Trimmomatic v0.36 (136). The strand-specific RNA-seq reads were then mapped to their respective genome using Hisat2 v2.1.0 (137) and a genome-guided transcriptome assembly was generated using default options in Trinity (138). A second transcript assembly was generated using StringTie2 v2.1.6 (139). Here, transcripts were assembled independently for each sample and a final set of unique transcripts was computed using the –merge function.

#### Gene prediction

Gene models were initially predicted separately for both *F. hygrometrica* accessions, using BRAKER2 v2.1.0 (122). The prediction algorithm was first trained in the GeneMark-ETP+ mode, providing mapped RNAseq data, proteome data of Viridiplantae species retrieved from OrthoDB_v10 (140), transcriptome data of *P. patens* retrieved from Phytozome13 (25, 141), and transcriptome assemblies of *F. hygrometrica* (see previous paragraph). The resulting species model was used in a second BRAKER2 run, omitting further training, and providing all available evidence to make a first comprehensive gene prediction and annotation. The evidence for this run included hint files generated from the repetitive element annotation by RepeatModeler2 v2.0 (113), RNAseq coverage data from all available samples, and *de novo* assembled transcripts produced with Trinity (138) and StringTie2 v2.1.6 (139) as described in the previous paragraph (Supplementary_Table_6).

#### Consolidating gene models of the two F. hygrometrica accessions

In an attempt to consolidate the previously generated gene predictions of the two *F. hygrometrica* accessions we employed the CGP extension of Augustus v3.3.1 (142) to transfer missing annotations between the two genomes. The resulting annotations, however, showed significantly worse BUSCO scores compared to the original annotations generated with the BRAKER2 pipeline. Visual inspection of the newly generated gene models showed that many previously well supported annotations were fragmented, leading to an overall decrease of annotation quality. Therefore, we decided to focus on gene models that were potentially missed during the gene prediction process in one or the other accession. To do so, we used the genBlastG algorithm v1.39 (123) to identify homologous regions of predicted gene sequences in the alternate accession’s genome and build valid exon-intron structures based on high-scoring sequence pairs and intrinsic sequence signals. We then tested if these newly generated gene models overlap with annotations generated by the BRAKER2 pipeline utilizing the -- intersect option of the BEDTools suite v2.26 (143). Only non-overlapping models generated by genBlastG were retained and added to the respective accession’s gene annotation file. To make these gene models distinguishable from the original annotation, the prefix “genb_” was added in front of the respective gene identifier (Supplementary_Table_6).

#### Filtering of incomplete gene models

All annotations were filtered to remove incomplete gene models before further analysis. Filtering was done using the program gFACs v1.1.2 (144). Models missing start or stop codons, showing incompleteness at their 5’ or 3’ ends or containing any in-frame stop codons were removed during filtering. Additionally, thresholds for minimum exon size (1), minimum intron size (10), and minimum total CDS size (225) were applied to remove short gene models.

### Whole genome alignments and collinearity analyses

We used dot plots generated with D-GENIES v1.2.0 (145) to assess collinearity between genomes of the two *F. hygrometrica* accessions as well as between *F. hygrometrica* and *P. patens* (25). We aligned the genomes using Minimap2 v2.17 (146). We excluded matches with less than 90% sequence identity when aligning genomes of the two *F. hygrometrica* accessions, while a threshold of 70% was used for alignments between *F. hygrometrica* and *P. patens*.

To assess structural variation between assemblies of the two *F. hygrometrica* accessions, we aligned them using the NUCmer module of the MUMmer package v4.0.0 (147). To visualise and classify the observed differences, we submitted the resulting .delta file to the Assemblytics web service v1.2.1 (148) (for further details see Supplementary Information).

We utilized the MCScan algorithm (149) and MCScanX toolkit (150) to assess collinearity within and between the studied genomes. We used peptide sequences of primary transcripts as input to an all-vs-all homology search with the BLAST algorithm (151), as recommended in the MCScanX documentation. The resulting tabular output was fed into the MCScan algorithm (149) to establish blocks of collinear genes using default parameters. We calculated synonymous and non-synonymous substitution rates for each syntenic gene pair using the tools supplied with the MCScanX toolkit (150). We visualized collinearity within genomes using circular plots generated with Circos v0.69-9 (152), while SynVisio (153) was employed to visualize collinearity between genomes.

### Gene set comparison of the *F. hygrometrica* and *P. patens* genomes

We created orthogroups, groups of genes descended from a common ancestor, using 38 plant proteomes including six species of green and streptophyte algae, 12 bryophytes, and 20 vascular plants representing all major lineages of land plants (Supplementary_Table_9, Supplementary_Table_10). OrthoFinder v.2.5.2 (154) analysis was run using default parameters. We obtained the species tree from orthogroup gene trees using the algorithm provided in OrthoFinder v.2.5.2. The species tree was converted into a time tree (ultrametric tree with branch length in time units) using the ete toolkit v3 (155). To infer gene family evolution on the branches leading to *F. hygrometrica* and *P. patens* from their common ancestor we used COUNT (156). COUNT applies a phylogenetic birth-and-death model to reconstruct the evolution of gene numbers in gene families along a phylogenetic tree taking into account the processes of gene loss, gene gain and duplication. All three parameters vary by the edges of the phylogenetic tree and by family, the latter according to a discretized gamma distribution. We used likelihood optimization to obtain numerical estimates for these parameters. To do so we performed model optimization in a model hierarchy starting with the simplest model and changing only one parameter at a time and retained parameters that led to the most significant improvement of the likelihood value. The final model included variable duplication rates, and edge length as well as duplication and loss rates varied according to a discrete gamma distribution with two parameters (length_k=2_dupl_k=2_loss_k=2). Gain was modelled with a simple gamma distribution as its inclusion did not influence the likelihood value significantly. Using the model parameter estimates, we calculated posterior probabilities for gene family expansion/contraction as well as gain/loss for each family running the posteriors module of COUNT (Supplementary_Table_11).

### Functional annotation of predicted genes

To obtain the functional annotation for the *F. hygrometrica* genes, we used two approaches. In particular, we assigned GO annotations to the gene models of *F. hygrometrica* using the eggNOG-mapper v2 (157) and InterProScan v5 (158) (Supplementray_Table_18-21.

## DECLARATIONS

### Ethics approval and consent to participate

Not applicable.

### Consent for publication

Not applicable.

### Availability of data and materials

Raw DNA and RNA sequencing data used in this publication was submitted to NCBI Short Read Archive (SRA) under the BioProject ID PRJNA816911 (SRA submission SUB11197892) and to the European Nucleotide Archive (ENA) under study accession number PRJEB36328 for the Zurich accession, respectively. For the UConn strain, raw DNA and RNA data were deposited at the CNGB data center (https://db.cngb.org/) under the project number CNP0002793. Genome assembly files, their annotations, and all supplementary tables will be deposited on figshare upon acceptance. A genome browser with Blast utilities will be made public upon acceptance.

### Competing interests

The authors declare that they have no competing interests.

### Funding

This study was financially supported by grants of the Swiss National Science Foundation (grant nos. 131726, 160004 and 184826 to PS), a pilot grant of the University Research Priority Program “Evolution in Action” of the University of Zurich to PS and AK, a Georges and Antoine Claraz Foundation grant to AK, MW, and PS, and a doctoral research grant (UZH Candoc, formerly Forschungskredit) to MW. This project was also carried out in the framework of MAdLand (https://madland.science, DFG priority programme 2237, PS-1111/1 to PS, RE 1697/19-1 and 20-1 to SAR). NR and BG were supported by a grant from the US National Science Foundation (DEB-1146295 & 1753811). The study was also funded by the Shenzhen Urban Management Bureau Fund (202005) to YL. The authors thank Yang Peng at the Shenzhen Fairy Lake Botanical Garden for laboratory assistance. This work was supported by China National GeneBank (CNGB; https://www.cngb.org/). RR received funding from the Deutsche Forschungsgemeinschaft DFG under CRC-Transregio 141 (project B02).

### Authors’ contributions

PS and YL conceptualized the study. PS, YL, SD, BG, NR, MW and SR generated primary sequence data. EMT carried out genome size measurements. PS and JY assembled the genomes. AK, YL, JY, NG, DL, LW and PS carried out genome annotation. AK and PS carried out detailed comparative genomic analyses. PS and AK wrote the manuscript. All co-authors revised and approved the final manuscript.

## Acknowledgements

We are grateful for Andrea Patrignani, Lucy Poveda, Weihong Qi, and Catharine Aquino for assistance in sequencing at the Functional Genomic Center Zurich (FGCZ). We are also thankful for the S3IT team and the ScienceCloud infrastructure at the University of Zurich for providing computational resources.

## Notes

### Competing Interest Statement

The authors have declared no competing interest.

## REFERENCES

1. Marks RA, Hotaling S, Frandsen PB, VanBuren R. Representation and participation across 20 years of plant genome sequencing. Nat plants [Internet]. 2021;7(12):1571–8. Available from: http://www.ncbi.nlm.nih.gov/pubmed/34845350

2. Kress WJ, Soltis DE, Kersey PJ, Wegrzyn JL, Leebens-Mack JH, Gostel MR, et al. Green plant genomes: What we know in an era of rapidly expanding opportunities. Proc Natl Acad Sci U S A [Internet]. 2022;119(4):1–9. Available from: http://www.ncbi.nlm.nih.gov/pubmed/35042803

3. Chen F, Dong W, Zhang J, Guo X, Chen J, Wang Z, et al. The sequenced angiosperm genomes and genome databases. Front Plant Sci. 2018;9(April):1–14.

4. Wendel JF, Jackson SA, Meyers BC, Wing RA. Evolution of plant genome architecture. Genome Biol [Internet]. 2016;17(1):1–14. Available from: http://dx.doi.org/10.1186/s13059-016-0908-1

5. Szövényi P, Gunadi A, Li F-W. Charting the genomic landscape of seed-free plants. Nat plants [Internet]. 2021;7(5):554–65. Available from: http://dx.doi.org/10.1038/s41477-021-00888-z

6. Hufford MB, Seetharam AS, Woodhouse MR, Chougule KM, Ou S, Liu J, et al. De novo assembly, annotation, and comparative analysis of 26 diverse maize genomes. Science (80-). 2021;373(6555):655–62.

7. Li H, Wang S, Chai S, Yang Z, Zhang Q, Xin H, et al. Graph-based pan-genome reveals structural and sequence variations related to agronomic traits and domestication in cucumber. Nat Commun [Internet]. 2022;13(1):682. Available from: http://www.ncbi.nlm.nih.gov/pubmed/35115520

8. Hoopes G, Meng X, Hamilton JP, Achakkagari SR, de Alves Freitas Guesdes F, Bolger ME, et al. Phased, chromosome-scale genome assemblies of tetraploid potato reveal a complex genome, transcriptome, and predicted proteome landscape underpinning genetic diversity. Mol Plant [Internet]. 2022 Jan;1–17. Available from: https://linkinghub.elsevier.com/retrieve/pii/S167420522200003X

9. Qiao Q, Edger PP, Xue L, Qiong L, Lu J, Zhang Y, et al. Evolutionary history and pan-genome dynamics of strawberry (Fragaria spp.). Proc Natl Acad Sci U S A [Internet]. 2021;118(45). Available from: http://www.ncbi.nlm.nih.gov/pubmed/34697247

10. Lovell JT, Bentley NB, Bhattarai G, Jenkins JW, Sreedasyam A, Alarcon Y, et al. Four chromosome scale genomes and a pan-genome annotation to accelerate pecan tree breeding. Nat Commun [Internet]. 2021;12(1):4125. Available from: http://dx.doi.org/10.1038/s41467-021-24328-w

11. Qin P, Lu H, Du H, Wang H, Chen W, Chen Z, et al. Pan-genome analysis of 33 genetically diverse rice accessions reveals hidden genomic variations. Cell [Internet]. 2021;184(13):3542–3558.e16. Available from: https://doi.org/10.1016/j.cell.2021.04.046

12. Tao Y, Luo H, Xu J, Cruickshank A, Zhao X, Teng F, et al. Extensive variation within the pan-genome of cultivated and wild sorghum. Nat plants [Internet]. 2021;7(6):766–73. Available from: http://dx.doi.org/10.1038/s41477-021-00925-x

13. Li J, Yuan D, Wang P, Wang Q, Sun M, Liu Z, et al. Cotton pan-genome retrieves the lost sequences and genes during domestication and selection. Genome Biol [Internet]. 2021;22(1):119. Available from: http://www.ncbi.nlm.nih.gov/pubmed/33892774

14. Hübner S, Bercovich N, Todesco M, Mandel JR, Odenheimer J, Ziegler E, et al. Sunflower pan-genome analysis shows that hybridization altered gene content and disease resistance. Nat plants [Internet]. 2019;5(1):54–62. Available from: http://dx.doi.org/10.1038/s41477-018-0329-0

15. Gao L, Gonda I, Sun H, Ma Q, Bao K, Tieman DM, et al. The tomato pan-genome uncovers new genes and a rare allele regulating fruit flavor. Nat Genet [Internet]. 2019;51(6):1044–51. Available from: http://dx.doi.org/10.1038/s41588-019-0410-2

16. Zhao Q, Feng Q, Lu H, Li Y, Wang A, Tian Q, et al. Pan-genome analysis highlights the extent of genomic variation in cultivated and wild rice. Nat Genet [Internet]. 2018;50(2):278–84. Available from: http://dx.doi.org/10.1038/s41588-018-0041-z

17. Wang W, Mauleon R, Hu Z, Chebotarov D, Tai S, Wu Z, et al. Genomic variation in 3,010 diverse accessions of Asian cultivated rice. Nature [Internet]. 2018;557(7703):43–9. Available from: http://www.ncbi.nlm.nih.gov/pubmed/29695866

18. Gordon SP, Contreras-Moreira B, Woods DP, Des Marais DL, Burgess D, Shu S, et al. Extensive gene content variation in the Brachypodium distachyon pan-genome correlates with population structure. Nat Commun [Internet]. 2017;8(1):2184. Available from: http://dx.doi.org/10.1038/s41467-017-02292-8

19. Golicz AA, Bayer PE, Barker GC, Edger PP, Kim H, Martinez PA, et al. The pangenome of an agronomically important crop plant Brassica oleracea. Nat Commun [Internet]. 2016;7:13390. Available from: http://dx.doi.org/10.1038/ncomms13390

20. Li Y, Zhou G, Ma J, Jiang W, Jin L, Zhang Z, et al. De novo assembly of soybean wild relatives for pan-genome analysis of diversity and agronomic traits. Nat Biotechnol [Internet]. 2014 Oct;32(10):1045–52. Available from: http://www.ncbi.nlm.nih.gov/pubmed/25218520

21. Rensing SA. Why we need more non-seed plant models. New Phytol [Internet]. 2017 Oct 13;216(2):355–60. Available from: https://onlinelibrary.wiley.com/doi/10.1111/nph.14464

22. Wickell D, Kuo L-Y, Yang H-P, Dhabalia Ashok A, Irisarri I, Dadras A, et al. Underwater CAM photosynthesis elucidated by Isoetes genome. Nat Commun [Internet]. 2021;12(1):6348. Available from: http://www.ncbi.nlm.nih.gov/pubmed/34732722

23. Carey SB, Jenkins J, Lovell JT, Maumus F, Sreedasyam A, Payton AC, et al. Gene-rich UV sex chromosomes harbor conserved regulators of sexual development. Sci Adv. 2021;7(27).

24. Silva AT, Gao B, Fisher KM, Mishler BD, Ekwealor JTB, Stark LR, et al. To dry perchance to live: Insights from the genome of the desiccation-tolerant biocrust moss Syntrichia caninervis. Plant J [Internet]. 2021 Mar;105(5):1339–56. Available from: http://www.ncbi.nlm.nih.gov/pubmed/33277766

25. Lang D, Ullrich KK, Murat F, Fuchs J, Jenkins J, Haas FB, et al. The Physcomitrella patens chromosome-scale assembly reveals moss genome structure and evolution. Plant J. 2018;93(3):515–33.

26. Yu J, Cai Y, Zhu Y, Zeng Y, Dong S, Zhang K, et al. Chromosome-Level Genome Assemblies of Two Hypnales (Mosses) Reveal High Intergeneric Synteny. Genome Biol Evol. 2022;14(2):1–6.

27. Li F-W, Nishiyama T, Waller M, Frangedakis E, Keller J, Li Z, et al. Anthoceros genomes illuminate the origin of land plants and the unique biology of hornworts. Nat Plants [Internet]. 2020 Mar 13;6(3):259–72. Available from: http://dx.doi.org/10.1038/s41477-020-0618-2

28. Wolf PG, Sessa EB, Marchant DB, Li FW, Rothfels CJ, Sigel EM, et al. An exploration into fern genome space. Genome Biol Evol. 2015;7(9):2533–44.

29. Marchant DB, Sessa EB, Wolf PG, Heo K, Barbazuk WB, Soltis PS, et al. The C-Fern (Ceratopteris richardii) genome: insights into plant genome evolution with the first partial homosporous fern genome assembly. Sci Rep [Internet]. 2019;9(1):18181. Available from: http://www.ncbi.nlm.nih.gov/pubmed/31796775

30. Rahmatpour N, Perera N V., Singh V, Wegrzyn JL, Goffinet B. High gene space divergence contrasts with frozen vegetative architecture in the moss family Funariaceae. Mol Phylogenet Evol [Internet]. 2021;154(January 2020):106965. Available from: https://doi.org/10.1016/j.ympev.2020.106965

31. Johnson MG, Malley C, Goffinet B, Shaw AJ, Wickett NJ. A phylotranscriptomic analysis of gene family expansion and evolution in the largest order of pleurocarpous mosses (Hypnales, Bryophyta). Mol Phylogenet Evol [Internet]. 2016 May;98:29–40. Available from: http://dx.doi.org/10.1016/j.ympev.2016.01.008

32. Buck WR, Shaw AJ, Goffinet B. Morphology, anatomy, and classification of the Bryophyta. In: Shaw AJ, editor. Bryophyte Biology [Internet]. Cambridge: Cambridge University Press; 2008. p. 55–138. Available from: https://www.cambridge.org/core/product/identifier/CBO9780511754807A009/type/book_part

33. Rensing SA, Goffinet B, Meyberg R, Wu SZ, Bezanilla M. The moss physcomitrium (Physcomitrella) patens: A model organism for non-seed plants. Plant Cell. 2020;32(5):1361– 76.

34. Medina R, Johnson MG, Liu Y, Wickett NJ, Shaw AJ, Goffinet B. Phylogenomic delineation of Physcomitrium (Bryophyta: Funariaceae) based on targeted sequencing of nuclear exons and their flanking regions rejects the retention of Physcomitrella, Physcomitridium and Aphanorrhegma. J Syst Evol. 2019;57(4):404–17.

35. Liu Y, Budke JM, Goffinet B. Phylogenetic inference rejects sporophyte based classification of the Funariaceae (Bryophyta): rapid radiation suggests rampant homoplasy in sporophyte evolution. Mol Phylogenet Evol [Internet]. 2012 Jan;62(1):130–45. Available from: http://dx.doi.org/10.1016/j.ympev.2011.09.010

36. Fernandez-Pozo N, Haas FB, Gould SB, Rensing SA. An overview of bioinformatics, genomics and transcriptomics resources for bryophytes. J Exp Bot [Internet]. 2022 Feb 11;54:1–54. Available from: http://www.ncbi.nlm.nih.gov/pubmed/35148385

37. Medina R, Johnson M, Liu Y, Wilding N, Hedderson TA, Wickett N, et al. Evolutionary dynamism in bryophytes: Phylogenomic inferences confirm rapid radiation in the moss family Funariaceae. Mol Phylogenet Evol [Internet]. 2018 Dec 5;120:240–7. Available from: https://doi.org/10.1016/j.ympev.2017.12.002

38. Beike AK, von Stackelberg M, Schallenberg-Rüdinger M, Hanke ST, Follo M, Quandt D, et al. Molecular evidence for convergent evolution and allopolyploid speciation within the Physcomitrium-Physcomitrella species complex. BMC Evol Biol [Internet]. 2014 Jul 11;14:158. Available from: http://www.ncbi.nlm.nih.gov/pubmed/25015729

39. Fife A. A generic revision of the Funariaceae (Bryophyta: Musci). Part I. J Hattori Bot Lab. 1985;58:149–196.

40. Budke JM, Goffinet B. Comparative Cuticle Development Reveals Taller Sporophytes Are Covered by Thicker Calyptra Cuticles in Mosses. Front Plant Sci [Internet]. 2016 Jun 14;7(JUNE2016):1–11. Available from: http://journal.frontiersin.org/Article/10.3389/fpls.2016.00832/abstract

41. McDaniel SF, Von Stackelberg M, Richardt S, Quatrano RS, Reski R, Rensing SA. The speciation history of the physcomitrium - Physcomitrella species complex. Evolution (N Y). 2010;64(1):217–31.

42. Reski R, Faust M, Wang X-H, Wehe M, Abel WO. Genome analysis of the moss Physcomitrella patens (Hedw.) B.S.G. Mol Gen Genet MGG [Internet]. 1994 Jul;244(4):352–9. Available from: https://www.libnauka.ru/item.php?doi=10.7868/S0132342317030071

43. Rice A, Glick L, Abadi S, Einhorn M, Kopelman NM, Salman-Minkov A, et al. The Chromosome Counts Database (CCDB) - a community resource of plant chromosome numbers. New Phytol [Internet]. 2015 Apr;206(1):19–26. Available from: http://www.ncbi.nlm.nih.gov/pubmed/25423910

44. Fritsch R. Index to Bryophyte Chromosome Counts. J. Cramer; 1991. (Bryophytorum bibliotheca).

45. Kapila S. Cytological observations on some west Himalayan mosses. Proc Indian Sci Congr Assoc. 1992;79:115–6.

46. Kapila S, Kumar SS. Cytological observations on some West Himalayan mosses. Hikobia. 1997;12:215–9.

47. Rensing SA, Lang D, Zimmer AD, Terry A, Salamov A, Shapiro H, et al. The Physcomitrella genome reveals evolutionary insights into the conquest of land by plants. Science (80-). 2008 Jan;319(5859):64–9.

48. Schween G, Gorr G, Hohe A, Reski R. Unique Tissue-Specific Cell Cycle in Physcomitrella. Plant Biol [Internet]. 2003 Jan;5(1):50–8. Available from: http://doi.wiley.com/10.1055/s-2003-37984

49. Voglmayr H. Nuclear DNA Amounts in Mosses (Musci). Ann Bot [Internet]. 2000 Apr;85(4):531–46. Available from: https://academic.oup.com/aob/article-lookup/doi/10.1006/anbo.1999.1103

50. Magdy M, Werner O, Mcdaniel SF, Goffinet B, Ros RM. Genomic scanning using AFLP to detect loci under selection in the moss Funaria hygrometrica along a climate gradient in the Sierra Nevada Mountains, Spain. Plant Biol. 2016;18(2):280–8.

51. Szövényi P, Devos N, Weston DJ, Yang X, Hock Z, Shaw JA, et al. Efficient Purging of Deleterious Mutations in Plants with Haploid Selfing. Genome Biol Evol [Internet]. 2014 May;6(5):1238–52. Available from: https://academic.oup.com/gbe/article-lookup/doi/10.1093/gbe/evu099

52. Szövényi P, Ullrich KK, Rensing SA, Lang D, van Gessel N, Stenøien HK, et al. Selfing in Haploid Plants and Efficacy of Selection: Codon Usage Bias in the Model Moss Physcomitrella patens. Genome Biol Evol [Internet]. 2017 Jun 1;9(6):1528–46. Available from: https://academic.oup.com/gbe/article-lookup/doi/10.1093/gbe/evx098

53. Haas FB, Fernandez-Pozo N, Meyberg R, Perroud PF, Göttig M, Stingl N, et al. Single Nucleotide Polymorphism Charting of P. patens Reveals Accumulation of Somatic Mutations During in vitro Culture on the Scale of Natural Variation by Selfing. Front Plant Sci. 2020;11(July):1–18.

54. Bengtsson BO, Cronberg N. The effective size of bryophyte populations. J Theor Biol. 2009;258(1):121–6.

55. Fritsch R. Index to bryophyte chromosome counts. Vol. 40, Bryophytorum Bibliotheca. 1991. 1–352 p.

56. Pellicer J, Leitch IJ. The Plant DNA C-values database (release 7.1): an updated online repository of plant genome size data for comparative studies. New Phytol [Internet]. 2020 Apr 8;226(2):301–5. Available from: https://onlinelibrary.wiley.com/doi/10.1111/nph.16261

57. Szövényi P, Perroud P-F, Symeonidi A, Stevenson S, Quatrano RS, Rensing SA, et al. De novo assembly and comparative analysis of the Ceratodon purpureus transcriptome. Mol Ecol Resour [Internet]. 2015 Jan;15(1):203–15. Available from: http://doi.wiley.com/10.1111/1755-0998.12284

58. Gao B, Chen MX, Li XS, Zhang DY, Wood AJ. Ancestral gene duplications in mosses characterized by integrated phylogenomic analyses. 2022;60(1):144–59.

59. Pellicer J, Hidalgo O, Dodsworth S, Leitch I. Genome Size Diversity and Its Impact on the Evolution of Land Plants. Genes (Basel) [Internet]. 2018 Feb 14;9(2):88. Available from: https://www.mdpi.com/2073-4425/9/2/88

60. Wang D, Zheng Z, Li Y, Hu H, Wang Z, Du X, et al. Which factors contribute most to genome size variation within angiosperms? Ecol Evol [Internet]. 2021 Mar 31;11(6):2660–8. Available from: https://onlinelibrary.wiley.com/doi/10.1002/ece3.7222

61. Suda J, Meyerson LA, Leitch IJ, Pyšek P. The hidden side of plant invasions: the role of genome size. New Phytol [Internet]. 2015 Feb;205(3):994–1007. Available from: http://www.ncbi.nlm.nih.gov/pubmed/25323486

62. Bilinski P, Albert PS, Berg JJ, Birchler JA, Grote MN, Lorant A, et al. Parallel altitudinal clines reveal trends in adaptive evolution of genome size in Zea mays. PLoS Genet [Internet]. 2018;14(5):e1007162. Available from: http://www.ncbi.nlm.nih.gov/pubmed/29746459

63. Bromham L, Hua X, Lanfear R, Cowman PF. Exploring the Relationships between Mutation Rates, Life History, Genome Size, Environment, and Species Richness in Flowering Plants. Am Nat [Internet]. 2015 Apr;185(4):507–24. Available from: https://www.journals.uchicago.edu/doi/10.1086/680052

64. Beric A, Mabry ME, Harkess AE, Brose J, Schranz ME, Conant GC, et al. Comparative phylogenetics of repetitive elements in a diverse order of flowering plants (Brassicales). G3 (Bethesda) [Internet]. 2021 May 16;11(7). Available from: http://www.ncbi.nlm.nih.gov/pubmed/33993297

65. Elliott TA, Gregory TR. What’s in a genome? The C-value enigma and the evolution of eukaryotic genome content. Philos Trans R Soc Lond B Biol Sci [Internet]. 2015 Sep 26;370(1678):20140331. Available from: http://www.ncbi.nlm.nih.gov/pubmed/26323762

66. Lisch D. How important are transposons for plant evolution? Nat Rev Genet [Internet]. 2013 Jan;14(1):49–61. Available from: http://www.ncbi.nlm.nih.gov/pubmed/23247435

67. Lysak MA, Koch MA, Beaulieu JM, Meister A, Leitch IJ. The dynamic ups and downs of genome size evolution in Brassicaceae. Mol Biol Evol [Internet]. 2009 Jan;26(1):85–98. Available from: http://www.ncbi.nlm.nih.gov/pubmed/18842687

68. Kingdom U. Heterochromatin diversity in two species of Pellia ( Hepaticae ) as revealed by C-, Q-, N- and Hoechst 33258-banding. 1985;(77):378–86.

69. Newton ME. Developmental Aspects of Chromatin Condensation in Liverworts Author ( s ): M . E . Newton Published by : American Bryological and Lichenological Society Stable URL : http://www.jstor.com/stable/3243101 Developmental Aspects of Chromatin Condensation in L. Bryologist. 1987;90(4):376–82.

70. Lynch M, Conery JS. The origins of genome complexity. Science [Internet]. 2003 Nov 21;302(5649):1401–4. Available from: http://www.ncbi.nlm.nih.gov/pubmed/14631042

71. Perroud PF, Cove DJ, Quatrano RS, Mcdaniel SF. An experimental method to facilitate the identification of hybrid sporophytes in the moss Physcomitrella patens using fluorescent tagged lines. New Phytol [Internet]. 2011;191(1):301–6. Available from: http://www.ncbi.nlm.nih.gov/pmc/articles/PMC3445409/pdf/nihms267253.pdf

72. McDaniel SF, Perroud P. Invited perspective: bryophytes as models for understanding the evolution of sexual systems. Bryologist [Internet]. 2012 Mar;115(1):1–11. Available from: http://www.bioone.org/doi/abs/10.1639/0007-2745-115.1.1

73. Szövényi P, Ullrich KK, Rensing SA, Lang D, van Gessel N, Stenøien HK, et al. Selfing in Haploid Plants and Efficacy of Selection: Codon Usage Bias in the Model Moss Physcomitrella patens. Genome Biol Evol. 2017;9(6).

74. Zhang G, Ge C, Xu P, Wang S, Cheng S, Han Y, et al. The reference genome of Miscanthus floridulus illuminates the evolution of Saccharinae. Nat plants [Internet]. 2021;7(5):608–18. Available from: http://dx.doi.org/10.1038/s41477-021-00908-y

75. Shi J, Ma X, Zhang J, Zhou Y, Liu M, Huang L, et al. Chromosome conformation capture resolved near complete genome assembly of broomcorn millet. Nat Commun [Internet]. 2019;10(1):464. Available from: http://dx.doi.org/10.1038/s41467-018-07876-6

76. Dvorak J, Wang L, Zhu T, Jorgensen CM, Deal KR, Dai X, et al. Structural variation and rates of genome evolution in the grass family seen through comparison of sequences of genomes greatly differing in size. Plant J [Internet]. 2018;95(3):487–503. Available from: http://www.ncbi.nlm.nih.gov/pubmed/29770515

77. Massa AN, Wanjugi H, Deal KR, O’Brien K, You FM, Maiti R, et al. Gene space dynamics during the evolution of Aegilops tauschii, Brachypodium distachyon, Oryza sativa, and Sorghum bicolor genomes. Mol Biol Evol [Internet]. 2011 Sep;28(9):2537–47. Available from: http://www.ncbi.nlm.nih.gov/pubmed/21470968

78. Huang K, Rieseberg LH. Frequency, Origins, and Evolutionary Role of Chromosomal Inversions in Plants. Front Plant Sci [Internet]. 2020 Mar 18;11(March):1–13. Available from: https://www.frontiersin.org/article/10.3389/fpls.2020.00296/full

79. Yang Z, Ge X, Yang Z, Qin W, Sun G, Wang Z, et al. Extensive intraspecific gene order and gene structural variations in upland cotton cultivars. Nat Commun [Internet]. 2019;10(1):2989. Available from: http://dx.doi.org/10.1038/s41467-019-10820-x

80. Yates TB, Feng K, Zhang J, Singan V, Jawdy SS, Ranjan P, et al. The Ancient Salicoid Genome Duplication Event: A Platform for Reconstruction of De Novo Gene Evolution in Populus trichocarpa. Genome Biol Evol [Internet]. 2021;13(9):1–14. Available from: http://www.ncbi.nlm.nih.gov/pubmed/34469536

81. Kirbis A, Waller M, Ricca M, Bont Z, Neubauer A, Goffinet B, et al. Transcriptional Landscapes of Divergent Sporophyte Development in Two Mosses, Physcomitrium (Physcomitrella) patens and Funaria hygrometrica. Front Plant Sci [Internet]. 2020 Jun 10;11:747. Available from: https://www.frontiersin.org/article/10.3389/fpls.2020.00747/full

82. Stein JC, Yu Y, Copetti D, Zwickl DJ, Zhang L, Zhang C, et al. Genomes of 13 domesticated and wild rice relatives highlight genetic conservation, turnover and innovation across the genus Oryza. Nat Genet [Internet]. 2018;50(2):285–96. Available from: http://dx.doi.org/10.1038/s41588-018-0040-0

83. Zhang L, Ren Y, Yang T, Li G, Chen J, Gschwend AR, et al. Rapid evolution of protein diversity by de novo origination in Oryza. Nat Ecol Evol [Internet]. 2019;3(4):679–90. Available from: http://dx.doi.org/10.1038/s41559-019-0822-5

84. Yang X, Jawdy S, Tschaplinski TJ, Tuskan GA. Genome-wide identification of lineage-specific genes in Arabidopsis, Oryza and Populus. Genomics. 2009;93(5):473–80.

85. Xu Y, Wu G, Hao B, Chen L, Deng X, Xu Q. Identification, characterization and expression analysis of lineage-specific genes within sweet orange (Citrus sinensis). BMC Genomics [Internet]. 2015;16(1):1–10. Available from: http://dx.doi.org/10.1186/s12864-015-2211-z

86. Graham MA, Silverstein KAT, Cannon SB, VandenBosch KA. Computational identification and characterization of novel genes from legumes. Plant Physiol [Internet]. 2004 Jul;135(3):1179–97. Available from: http://www.ncbi.nlm.nih.gov/pubmed/15266052

87. Donoghue MT, Keshavaiah C, Swamidatta SH, Spillane C. Evolutionary origins of Brassicaceae specific genes in Arabidopsis thaliana. BMC Evol Biol [Internet]. 2011 Feb 18;11(1):47. Available from: http://www.ncbi.nlm.nih.gov/pubmed/21332978

88. Wang M, Li J, Wang P, Liu F, Liu Z, Zhao G, et al. Comparative Genome Analyses Highlight Transposon-Mediated Genome Expansion and the Evolutionary Architecture of 3D Genomic Folding in Cotton. Mol Biol Evol [Internet]. 2021 Aug 23;38(9):3621–36. Available from: http://www.ncbi.nlm.nih.gov/pubmed/33973633

89. Kou Y, Liao Y, Toivainen T, Lv Y, Tian X, Emerson JJ, et al. Evolutionary Genomics of Structural Variation in Asian Rice (Oryza sativa) Domestication. Mol Biol Evol [Internet]. 2020;37(12):3507–24. Available from: http://www.ncbi.nlm.nih.gov/pubmed/32681796

90. Guan Y, Liu L, Wang Q, Zhao J, Li P, Hu J, et al. Gene refashioning through innovative shifting of reading frames in mosses. Nat Commun [Internet]. 2018;9(1). Available from: http://dx.doi.org/10.1038/s41467-018-04025-x

91. Tautz D, Domazet-Lošo T. The evolutionary origin of orphan genes. Nat Rev Genet. 2011;12(10):692–702.

92. Bornberg-Bauer E, Hlouchova K, Lange A. Structure and function of naturally evolved de novo proteins. Curr Opin Struct Biol. 2021;68:175–83.

93. Rödelsperger C, Prabh N, Sommer RJ. New Gene Origin and Deep Taxon Phylogenomics: Opportunities and Challenges. Trends Genet [Internet]. 2019 Dec 1;35(12):914–22. Available from: https://doi.org/10.1016/j.tig.2019.08.007

94. Quatrano RS, McDaniel SF, Khandelwal A, Perroud PF, Cove DJ. Physcomitrella patens: mosses enter the genomic age. Curr Opin Plant Biol. 2007;10(2):182–9.

95. Bowman JL, Kohchi T, Yamato KT, Jenkins J, Shu S, Ishizaki K, et al. Insights into Land Plant Evolution Garnered from the Marchantia polymorpha Genome. Cell [Internet]. 2017 Oct 5;171(2):287–304.e15. Available from: http://www.ncbi.nlm.nih.gov/pubmed/28985561

96. Carey SB, Jenkins J, Lovell JT, Maumus F, Sreedasyam A, Payton AC, et al. The Ceratodon purpureus genome uncovers structurally complex, gene rich sex chromosomes. bioRxiv. 2020 Dec;2020.07.03.163634.

97. Montgomery SA, Tanizawa Y, Galik B, Wang N, Ito T, Mochizuki T, et al. Chromatin Organization in Early Land Plants Reveals an Ancestral Association between H3K27me3, Transposons, and Constitutive Heterochromatin. Curr Biol. 2020;30(4):573–588.e7.

98. Bopp M. Die Entwicklung von Zelle und Kern im Protonema von Funaria hygrometrica Sibth. Planta [Internet]. 1955 Jul;45(6):573–90. Available from: http://link.springer.com/10.1007/BF01911551

99. Diop SI, Subotic O, Giraldo-Fonseca A, Waller M, Kirbis A, Neubauer A, et al. A pseudomolecule-scale genome assembly of the liverwort Marchantia polymorpha. Plant J. 2020;101(6).

100. Wyatt R, Stoneburner A. Biosystematics of Bryophytes: An Overview. In: Grant WFBT-PB, editor. Plant Biosystematics [Internet]. Elsevier; 1984. p. 519–42. Available from: https://www.sciencedirect.com/science/article/pii/B9780122956805500366

101. Porebski S, Bailey LG, Baum BR. Modification of a CTAB DNA extraction protocol for plants containing high polysaccharide and polyphenol components. Plant Mol Biol Report. 1997;15(1):8–15.

102. Xie T, Zheng JF, Liu S, Peng C, Zhou YM, Yang QY, et al. De novo plant genome assembly based on chromatin interactions: A case study of arabidopsis thaliana. Mol Plant. 2015;8(3):489–92.

103. Koren S, Walenz BP, Berlin K, Miller JR, Bergman NH, Phillippy AM. Canu: Scalable and accurate long-read assembly via adaptive κ-mer weighting and repeat separation. Genome Res. 2017 May;27(5):722–36.

104. Putnam NH, O’Connell BL, Stites JC, Rice BJ, Blanchette M, Calef R, et al. Chromosome-scale shotgun assembly using an in vitro method for long-range linkage. Genome Res. 2016 Mar;26(3):342–50.

105. Durand NC, Shamim MS, Machol I, Rao SSP, Huntley MH, Lander ES, et al. Juicer Provides a One-Click System for Analyzing Loop-Resolution Hi-C Experiments. Cell Syst. 2016 Jul;3(1):95– 8.

106. Dudchenko O, Batra SS, Omer AD, Nyquist SK, Hoeger M, Durand NC, et al. De novo assembly of the Aedes aegypti genome using Hi-C yields chromosome-length scaffolds. Science (80-). 2017 Apr;356(6333):92–5.

107. Durand NC, Robinson JT, Shamim MS, Machol I, Mesirov JP, Lander ES, et al. Juicebox Provides a Visualization System for Hi-C Contact Maps with Unlimited Zoom. Cell Syst. 2016 Jul;3(1):99– 101.

108. Li H. Aligning sequence reads, clone sequences and assembly contigs with BWA-MEM. arXiv Prepr arXiv. 2013;00(00):3.

109. Walker BJ, Abeel T, Shea T, Priest M, Abouelliel A, Sakthikumar S, et al. Pilon: An Integrated Tool for Comprehensive Microbial Variant Detection and Genome Assembly Improvement. Wang J, editor. PLoS One [Internet]. 2014 Nov 19;9(11):e112963. Available from: https://dx.plos.org/10.1371/journal.pone.0112963

110. English AC, Richards S, Han Y, Wang M, Vee V, Qu J, et al. Mind the Gap : Upgrading Genomes with Pacific Biosciences RS Long-Read Sequencing Technology. 2012;7(11):1–12.

111. Vaser R, Sović I, Nagarajan N, Šikić M. Fast and accurate de novo genome assembly from long uncorrected reads. Genome Res [Internet]. 2017;27(5):737–46. Available from: http://www.ncbi.nlm.nih.gov/pubmed/28100585

112. Laetsch DR, Blaxter ML. BlobTools: Interrogation of genome assemblies. F1000Research. 2017 Jul;6:1287.

113. Flynn JM, Hubley R, Goubert C, Rosen J, Clark AG, Feschotte C, et al. RepeatModeler2 for automated genomic discovery of transposable element families. Proc Natl Acad Sci U S A. 2020 Apr;117(17):9451–7.

114. Price AL, Jones NC, Pevzner PA. De novo identification of repeat families in large genomes. Bioinformatics. 2005 Jun;21(SUPPL. 1).

115. Bao Z, Eddy SR. Automated de novo identification of repeat sequence families in sequenced genomes. Genome Res. 2002 Aug;12(8):1269–76.

116. Ellinghaus D, Kurtz S, Willhoeft U. LTRharvest, an efficient and flexible software for de novo detection of LTR retrotransposons. BMC Bioinformatics. 2008 Jan;9.

117. Gremme G, Steinbiss S, Kurtz S. Genome tools: A comprehensive software library for efficient processing of structured genome annotations. IEEE/ACM Trans Comput Biol Bioinforma. 2013 May;10(3):645–56.

118. Ou S, Jiang N. LTR_retriever: A highly accurate and sensitive program for identification of long terminal repeat retrotransposons. Plant Physiol. 2018 Feb;176(2):1410–22.

119. Storer J, Hubley R, Rosen J, Wheeler TJ, Smit AF. The Dfam community resource of transposable element families, sequence models, and genome annotations. Mob DNA. 2021 Dec;12(1):2.

120. Smit A, Hubley R, Green P. RepeatMasker Open-4.0. 2015. p. <http://www.repeatmasker.org>.

121. Ou S, Su W, Liao Y, Chougule K, Agda JRA, Hellinga AJ, et al. Benchmarking transposable element annotation methods for creation of a streamlined, comprehensive pipeline. Genome Biol. 2019 Dec;20(1):275.

122. Brůna T, Hoff KJ, Lomsadze A, Stanke M, Borodovsky M. BRAKER2: automatic eukaryotic genome annotation with GeneMark-EP+ and AUGUSTUS supported by a protein database. NAR Genomics Bioinforma [Internet]. 2021 Jan 6;3(1):1–11. Available from: https://academic.oup.com/nargab/article/doi/10.1093/nargab/lqaa108/6066535

123. She R, Chu JS-C, Uyar B, Wang J, Wang K, Chen N. genBlastG: using BLAST searches to build homologous gene models. Bioinformatics. 2011 Aug;27(15):2141–3.

124. Llorens C, Futami R, Covelli L, Domínguez-Escribá L, Viu JM, Tamarit D, et al. The Gypsy Database (GyDB) of Mobile Genetic Elements: Release 2.0. Nucleic Acids Res. 2011 Jan;39(SUPPL. 1):D70–4.

125. Potter SC, Luciani A, Eddy SR, Park Y, Lopez R, Finn RD. HMMER web server: 2018 update. Nucleic Acids Res. 2018 Jul;46(W1):W200–4.

126. Eddy SR. Accelerated profile HMM searches. PLoS Comput Biol. 2011 Oct;7(10):1002195.

127. Edgar RC. MUSCLE: A multiple sequence alignment method with reduced time and space complexity. BMC Bioinformatics. 2004 Aug;5(1):113.

128. Saitou N, Nei M. The neighbor-joining method: a new method for reconstructing phylogenetic trees. Mol Biol Evol. 1987 Jul;4(4):406–25.

129. Kumar S, Stecher G, Li M, Knyaz C, Tamura K. MEGA X: Molecular Evolutionary Genetics Analysis across Computing Platforms. Battistuzzi FU, editor. Mol Biol Evol. 2018 Jun;35(6):1547–9.

130. Felsenstein J. CONFIDENCE LIMITS ON PHYLOGENIES: AN APPROACH USING THE BOOTSTRAP. Evolution (N Y). 1985 Jul;39(4):783–91.

131. Perlman PS, Boeke JD. Ring around the Retroelement. Vol. 303, Science. American Association for the Advancement of Science; 2004. p. 182–4.

132. Kimura M. A simple method for estimating evolutionary rates of base substitutions through comparative studies of nucleotide sequences. J Mol Evol. 1980 Jun;16(2):111–20.

133. Cock PJA, Antao T, Chang JT, Chapman BA, Cox CJ, Dalke A, et al. Biopython: Freely available Python tools for computational molecular biology and bioinformatics. Bioinformatics. 2009 Jun;25(11):1422–3.

134. Rensing S a, Ick J, Fawcett J a, Lang D, Zimmer A, Van de Peer Y, et al. An ancient genome duplication contributed to the abundance of metabolic genes in the moss Physcomitrella patens. BMC Evol Biol. 2007;7:130.

135. Wickham H. ggplot2: Elegant Graphics for Data Analysis. Springer-Verlag New York; 2016.

136. Bolger AM, Lohse M, Usadel B. Trimmomatic: A flexible trimmer for Illumina sequence data. Bioinformatics. 2014;30(15):2114–20.

137. Kim D, Langmead B, Salzberg SL. HISAT: A fast spliced aligner with low memory requirements. Nat Methods. 2015;12(4):357–60.

138. Grabherr MG, Haas BJ, Yassour M, Levin JZ, Thompson DA, Amit I, et al. Full-length transcriptome assembly from RNA-Seq data without a reference genome. Nat Biotechnol. 2011 Jul;29(7):644–52.

139. Kovaka S, Zimin A V., Pertea GM, Razaghi R, Salzberg SL, Pertea M. Transcriptome assembly from long-read RNA-seq alignments with StringTie2. Genome Biol. 2019 Dec;20(1):278.

140. Kriventseva E V., Kuznetsov D, Tegenfeldt F, Manni M, Dias R, Simão FA, et al. OrthoDB v10: Sampling the diversity of animal, plant, fungal, protist, bacterial and viral genomes for evolutionary and functional annotations of orthologs. Nucleic Acids Res. 2019 Jan;47(D1):D807–11.

141. Goodstein DM, Shu S, Howson R, Neupane R, Hayes RD, Fazo J, et al. Phytozome: A comparative platform for green plant genomics. Nucleic Acids Res [Internet]. 2012;40(D1):1178–86. Available from: http://www.ncbi.nlm.nih.gov/pmc/articles/PMC3245001/pdf/gkr944.pdf

142. König S, Romoth LW, Gerischer L, Stanke M. Simultaneous gene finding in multiple genomes. Bioinformatics. 2016 Jul;32(22):btw494.

143. Quinlan AR, Hall IM. BEDTools: A flexible suite of utilities for comparing genomic features. Bioinformatics. 2010;26(6):841–2.

144. Caballero M, Wegrzyn J. gFACs: Gene Filtering, Analysis, and Conversion to Unify Genome Annotations Across Alignment and Gene Prediction Frameworks. Genomics, Proteomics Bioinforma. 2019 Jun;17(3):305–10.

145. Cabanettes F, Klopp C. D-GENIES: Dot plot large genomes in an interactive, efficient and simple way. PeerJ. 2018 Jun;2018(6):e4958.

146. Li H. Minimap2: pairwise alignment for nucleotide sequences. Birol I, editor. Bioinformatics. 2018 Sep;34(18):3094–100.

147. Kurtz S, Phillippy A, Delcher AL, Smoot M, Shumway M, Antonescu C, et al. Versatile and open software for comparing large genomes. Genome Biol. 2004 Jan;5(2):R12.

148. Nattestad M, Schatz MC. Assemblytics: A web analytics tool for the detection of variants from an assembly. Bioinformatics. 2016;32(19):3021–3.

149. Tang H, Wang X, Bowers JE, Ming R, Alam M, Paterson AH. Unraveling ancient hexaploidy through multiply-aligned angiosperm gene maps. Genome Res. 2008 Dec;18(12):1944–54.

150. Wang Y, Tang H, Debarry JD, Tan X, Li J, Wang X, et al. MCScanX: A toolkit for detection and evolutionary analysis of gene synteny and collinearity. Nucleic Acids Res. 2012 Apr;40(7):e49.

151. Altschul SF, Gish W, Miller W, Myers EW, Lipman DJ. Basic local alignment search tool. J Mol Biol. 1990 Oct;215(3):403–10.

152. Krzywinski M, Schein J, Birol I, Connors J, Gascoyne R, Horsman D, et al. Circos: An information aesthetic for comparative genomics. Genome Res. 2009 Sep;19(9):1639–45.

153. Bandi V, Gutwin C. Interactive Exploration of Genomic Conversation. In: Proceedings of Graphics Interface 2020. Canadian Human-Computer Communications Society; 2020. p. 74– 83.

154. Emms DM, Kelly S. OrthoFinder: phylogenetic orthology inference for comparative genomics. Genome Biol [Internet]. 2019;20(1):238. Available from: http://www.ncbi.nlm.nih.gov/pubmed/31727128

155. Huerta-Cepas J, Serra F, Bork P. ETE 3: Reconstruction, Analysis, and Visualization of Phylogenomic Data. Mol Biol Evol [Internet]. 2016;33(6):1635–8. Available from: http://www.ncbi.nlm.nih.gov/pubmed/26921390

156. Csurös M. Count: evolutionary analysis of phylogenetic profiles with parsimony and likelihood. Bioinformatics [Internet]. 2010 Aug 1;26(15):1910–2. Available from: http://www.ncbi.nlm.nih.gov/pubmed/20551134

157. Cantalapiedra CP, Hernández-Plaza A, Letunic I, Bork P, Huerta-Cepas J. eggNOG-mapper v2: Functional Annotation, Orthology Assignments, and Domain Prediction at the Metagenomic Scale. Mol Biol Evol [Internet]. 2021 Dec 9;38(12):5825–9. Available from: http://www.ncbi.nlm.nih.gov/pubmed/34597405

158. Jones P, Binns D, Chang HY, Fraser M, Li W, McAnulla C, et al. InterProScan 5: Genome-scale protein function classification. Bioinformatics. 2014;30(9):1236–40.

