## Supplementary Information for "Genome dynamics in mosses: Extensive synteny coexists with a highly dynamic gene space"


1. **Genome size estimates**
2. **Manual revision of the assemblies**
3. **Whole genome alignment of the two *Funaria* accessions**
4. **Whole genome alignment of the *P. patens* and *F. hygrometrica* genomes**
5. **Gene set comparison of the *F. hygrometrica* accessions (Zurich and Uconn)**
6. **Phylogenetic analysis of LTR Copia and Gypsy super-families**
7. **References**

### 1.Genome size estimates

#### 1.1 Quantitative cell nuclei staining (FCM and FDM)

##### 1.1.1 *P. patens*, *F. hygrometrica* Zurich and Uconn accessions

*Plant Material*

Three gametophytic in vitro culture samples of *Funaria hygrometrica* Hedw.*,* (Funariaceae) Zurich*, Funaria hygrometrica* Hedw.*,* (Funariaceae) Uconn, and three *Physcomitrella patens* Gransden (Hedw.) Bruch & Schimp (Funariaceae) samples were analyzed for their genome size, or ploidy level, respectively. Plants were grown on BCD medium solidified with plant agar in 9 cm diameter petri dishes under continuous light (approx. 100 mikroE, 24 C°) for 1-2 month (Cove et al. 2009).

*Flow Cytometry (FCM) methods*

The gametophytes of *Physcomitrella patens* and *Funaria hygrometrica* were co-chopped according to the method of Galbraith et al. (Galbraith et al. 1983) together with *Solanum pseudocapsicum* as an internal standard (1.295 pg/1C, (Temsch, Greilhuber, and Krisai 2010)) in Otto´s buffer I (Otto et al. 1981). The resulting suspension was filtered through a 30 µm nylon mesh and treated with RNase A (0.15 mg/ml) for 30 min at 37°C in the water bath. Subsequently, the nuclei isolate was stained in propidium iodide (PI, 50 mg/l) supplemented Otto´s buffer II (Otto et al. 1981) and kept in the refrigerator until measurement on a CyFlow ML flow cytometer (Partec, Muenster, Germany), equipped with a green laser (100 mW, 532 nm, Cobolt Samba, Cobolt, Stockholm, Sweden). In total, 15,000 particles were measured for genome size measurement in the reference samples (samples 1 to 5), or 5,000 particles in the samples in question, respectively. The C-value was calculated according to the formula $1C_{obj}$ = (mean $G_2$ or $G_1$ nuclei fluorescence intensity peak $_{obj\ gam}$ / mean G1 nuclei fluorescence intensity peak $_{std\ spor}$) * 1 or 2C-value $_{std}$.

*Feulgen Densitometry (FDM) methods*

Our quantitative standard Feulgen procedure followed (Greilhuber and Temsch 2001). Gametophytes were fixated in methanol : acetic acid (3:1) and stored in 96% ethanol until further processing. Before staining, the fixated plantlets were washed together with the standard primary root tips (*Pisum sativum* ´Kleine Rheinländerin´, 1C=4.42pg, (Greilhuber and Ebert 1994)) in distilled water, hydrolyzed in 5N HCl for 60 minutes at 20°C (ultra-thermostate water bath). HCl was removed by few fast washes with distilled water and Schiff´s reagent was added and incubated over-night in the refrigerator. Several washes with $SO_2$-water were performed in order to remove the dye completely. All tissue was softened with 45% acetic acid and squashed on slides, covered with cover slips and frozen on a cold plate. After removal of the cover slips, the preparations were fixated in ethanol, dried and the integrated optical density (IOD) was measured in selected nuclei (telophases and/or prophases) using the Cell Image Retrieval and Evaluation System (CIRES, Kontron, Munich). The C-values were calculated similarly to the FCM results using the IODs of the standard as well as the object nuclei.

*Results*

Only one ploidy level was found in the isolates, haploid/diploid depending on either a main cell cycle arrest in $G_1$, or $G_2$ phases, respectively. In former approaches, in vitro cultures of *Physcomitrella patens* had shown a main cell cycle arrest in $G_2$ in contrast to the $G_1$ arrest common to wild plants (Reski et al. 1994). Since a valid result of C-value measurement by FCM bases on the exact knowledge of the cell cycle status of the main nuclei fraction, this ambiguity was to be clarified by means of mitotic nuclei measurement with Feulgen densitometry.

Considering the Feulgen evaluation of cell cycle status ($G_2$ arrest) at FCM measurement, the lower and higher ploidy level found in *Physcomitrium* (*Physcomitrella) patens* exhibits a mean 1C-value of 0.5382 pg, and 1.0854 pg, respectively. This is a 1:2.02 ratio. In *Funaria hygrometrica* Zurich accession, the lower 1C-value was 0.4084 pg, measured from two samples, and the higher was 0.8192 pg only from one sample available. This is a 1:2.01 ratio. Former approaches point to a 1Cx-value of 0.53 pg (Schween et al. 2003) in *P. patens* at a basic chromosome number of n = x = 27 (Reski et al. 1994), which differs only 1.5% from our nuclei DNA content measurement.

In *F. hygrometrica*, our estimate suggests a smaller genome size. Unfortunately, mitoses were not present in *F. hygrometrica* FDM preparations at all. Therefore, the classification of the interphase nuclei into $G_1$ or $G_2$ arrest classes is ambiguous. Schween et al. (2003) describes a phytohormone effect on the cell cycle arrest common to axenic *P. patens* cultures leading to cell cycle arrest in $G_2$ (2C) rather than $G_1$ (1C). Assuming that these findings are transferable to *F. hygrometrica*, its 1C-value equals 0.4084 pg. This corresponds to an approximately 399.4 Mbp genome size of the *F. hygrometrica* accession and about 526.4 Mpb genome size for *P. patens* using the experimental conversion factor: Genome size (bp) = $(0.978 \times 10^9) \times$ DNA content (pg) (Dolezel et al. 2003). Therefore, our estimates suggests that the *F. hygrometrica* genome is about 100-127 Mbp smaller than the *P. patens* genome. The most recent pseudochromosome-scale assembly of *P. patens* spans

462.3 Mbp of the estimated 518 Mbp genome size obtained by FCM (Lang et al. 2018). If a similar relationship between flow cytometry-based genome size estimate and actual assembly length applies to *F. hygrometrica*, its assembly length is expected to be about 357 Mbp.

The genome size of *F. hygrometrica* Uconn accession was estimated in the very same way using FCM, and by comparing *F. hygrometrica* (Uconn accession) with the standard material of *Oryza sativa* L. ssp. *japonica* (430 Mb, (Yu et al. 2002)). Two replicates were performed and the average genome size of *F. hygrometrica* (Uconn accession) was estimated as 370 Mb.

##### 1.2 K-mer-based genome size estimates

We carried out k-mer analysis using Illumina reads to obtain genome size estimates for the two *F. hygrometrica* accessions. We first run Jellyfish v2.3.0 (Marçais and Kingsford 2011) with a default k-mer value of 21 (jellyfish count -C -m 21 -s 1000000000 -t 28 *fastq -o reads.jf). We then converted the k-mer counts into histograms (jellyfish histo -t 28 reads.jf 1> reads.histo) and used Genomescope2 (Ranallo-Benavidez, Jaron, and Schatz 2020) with a “Max kmer coverage=1000000” cutoff to obtain genome size estimates (http://qb.cshl.edu/genomescope/). Our analyses resulted in a genome size estimate of 306,879,850-307,778,776 bp (minimum and maximum estimates) for the Zurich and 298,207,941-298,324,211 bp (min and max estimates) for the Uconn accession. Genomescope plots and estimates are reported below (Figure 1 and Table 1).

**A**

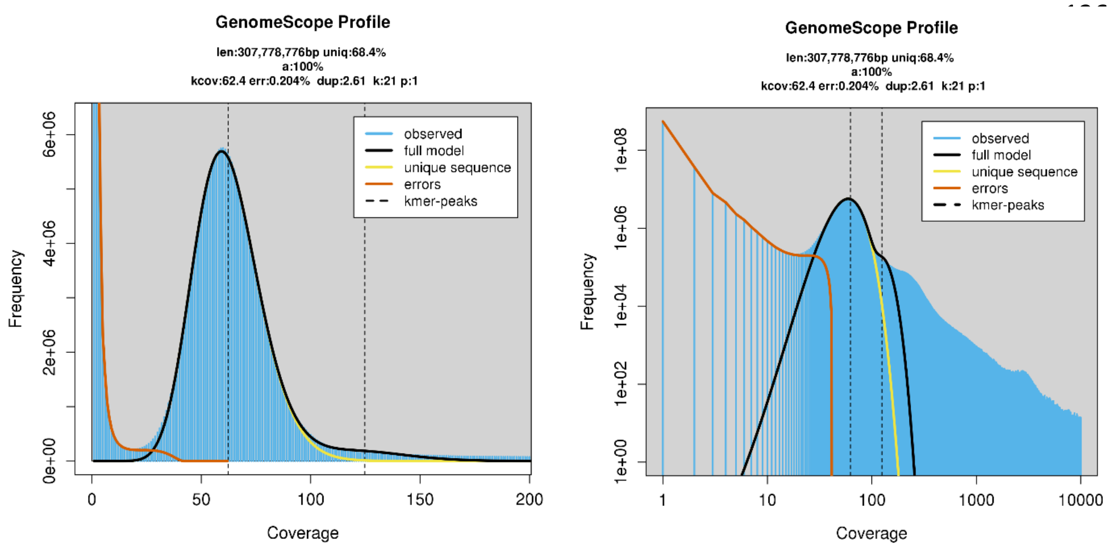


**B**

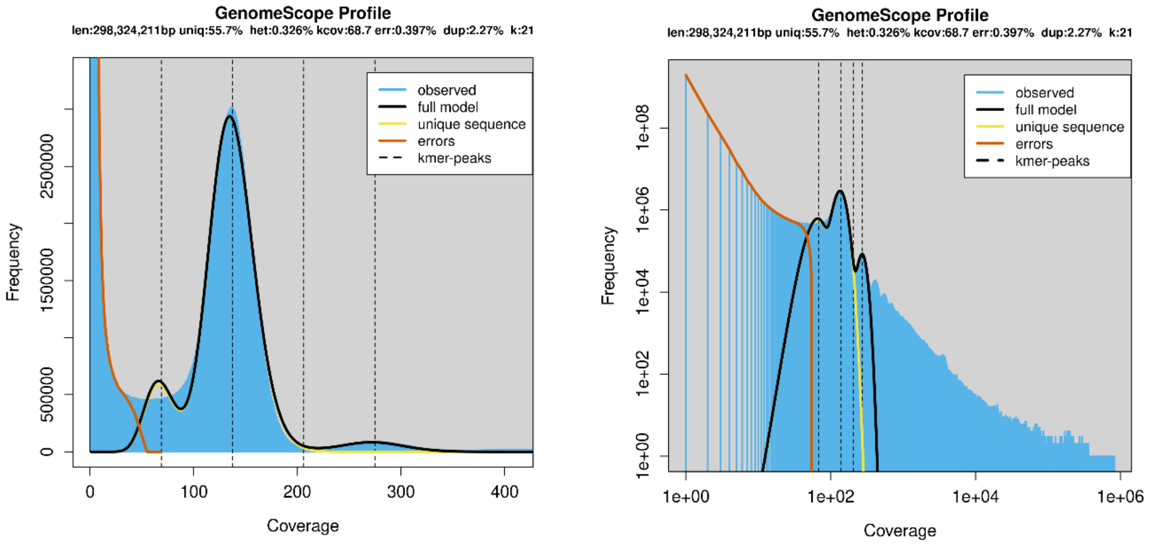


**Figure 1** Genomescope plots for the *F. hygrometrica* Zurich (A) and UConn accessions (B).

***F. hygrometrica* Zurich accession**

| property | min | max |
|---|---|---|
| Homozygous (a) | 100% | 100% |
| Genome Haploid Length | 306,879,850 bp | 307,778,776 bp |
| Genome Repeat Length | 96,833,278 bp | 97,116,926 bp |
| Genome Unique Length | 210,046,572 bp | 210,661,850 bp |
| Model Fit | 71.70% | 95.12% |
| Read Error Rate | 0.20% | 0.20% |

***F. hygrometrica* UConn accession**

| property | min | max |
|---|---|---|
| Heterozygosity | 0.02% | 0.03% |
| Genome Haploid Length | 298,207,941 bp | 298,324,211 bp |
| Genome Repeat Length | 132,220,786 bp | 132,272,339 bp |
| Genome Unique Length | 165,987,155 bp | 166,051,873 bp |
| Model Fit | 88.52% | 91.66% |
| Read Error Rate | 0.40% | 0.40% |

**Table 1** Genome size estimates obtained by Genomescope for the *F. hygrometrica* Zurich and UConn accessions.

#### 2. Manual revision of the assemblies

We assembled the Zurich accession`s genome using PacBio reads which were further scaffolded up using Chicago and Hi-C libraries. By contrast, the initial assembly of the Uconn accession was generated using Oxford-nanopore reads which contiguity was further improved using only Hi-C data. Details of the genome assemblies are described in the main text (methods).

We manually reviewed the accuracy of both final assemblies using the juicer/juicebox tools and to correct potential assembly errors. To do so, we mapped both the Chicago and Hi-C data (Zurich accession) or the Hi-C data (Uconn accession) onto the final genome assemblies and created Hi-C contact maps using the juicer/juicebox tools (Durand et al. 2016). We visually assessed the contact maps for potential errors including inversions, false joins/splits and falsely collapsed genomic regions. Neither the Chicago (Zurich accession) nor the Hi-C (Zurich and Uconn accessions) contact maps indicated the presence of large-scale assembly errors. Nevertheless, we discovered that two chromosome-scale scaffolds were falsely joined into a large scaffold in the Zurich accession`s initial assembly. This was clearly indicated by a markedly decreased contact density both in the Chicago and Hi-C libraries. To correct this error, we manually split this scaffold into two sub scaffolds. These steps are indicated on the figure below. We made no major editing on the scaffolds of the Uconn assembly.

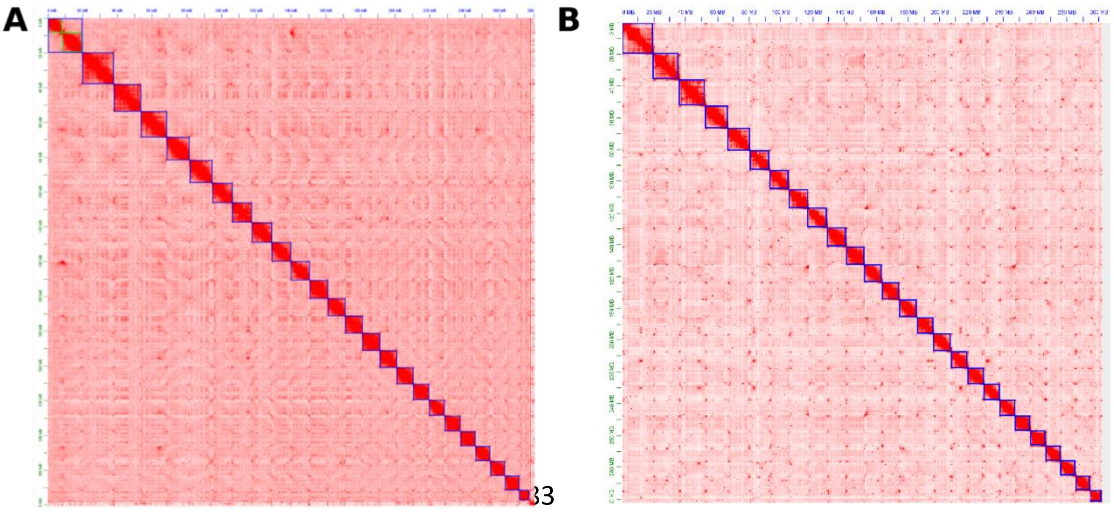


**Figure 2** Visualization of the revision made on the genome assembly of *F. hygrometrica* (Zurich accession). Left panel: Heatmap of Hi-C contacts within the initial

(unrevised) genome assembly. Blue rectangles represent 25 chromosome-scale scaffolds established in the initial assembly. We revised the assembly after reviewing Hi-C data, and split the largest scaffold (framed with a blue rectangle) into two chromosomal scaffolds (framed with green color). Right panel: Hi-C contact map of the Uconn accession.

##### 3. Whole genome alignment of the two *Funaria* accessions

We first aligned the two *F. hygrometrica* genomes (Zurich vs. Uconn) using default options with minimap2 (Li 2021) to evaluate their overall sequence divergence. We found that the genome of the two accessions were very similar. More specifically, only 7% and 15% of the genome sequences were specific (showed no similarity to the other genome at a similarity threshold of 50% at the nucleotide level) either to the Zurich or the Uconn accessions, respectively. Furthermore, sequence identity in about half of the genome was greater than 75% (Supplementary_Table_5). We found that overall sequence divergence across all alignments was on average 7.602% (median: 6.885 %, inter-quatile range [IQR:] 3.365- 9.865%) between the two strains.

To quantify the structural divergence between the genomes of the two *F. hygrometrica* accessions, we aligned them with mummer and used the Assemblytic (http://assemblytics.com/) server for visualization and classification of the differences observed. To verify these findings, we also created a whole genome alignment between the two *F. hygrometrica* accessions genomes using progressive Cactus v.2.0.5 with default options (Armstrong et al. 2020) and extracted SNPs and structural variants using haltools v.2.2 (Hickey et al. 2013). Despite being highly collinear, assembly length of the two accessions was different, the Uconn accession`s genome being 34 Mbp longer. This was partially due to structural variations between the two accessions` genome. We detected over 5700 (5760) structural variations in the Uconn compared to the Zurich assembly affecting in total over 14Mbp of the genome (Supplementary_Table_4). Repeat expansions/contractions (61%) were the most frequent structural variations which were followed by insertion/deletions (37%) while very few tandem expansions/contractions (2%) were observed. Insertions/deletions were mainly small-scale (50-500bp) while repeat expansions were dominated by large-scale (500-10kbp) variations (Figure 3). We also tabulated the number of inversions using the Cactus whole-genome alignment and haltools and by reactivating a currently uncommented portion of the Assemblytic pipeline script. We found that the absolute number of inversions detected by both alogorithms were very low about 1.1% (59). Therefore, inversion events are one of the least frequent structural variants between the two *Funaria* accessions.

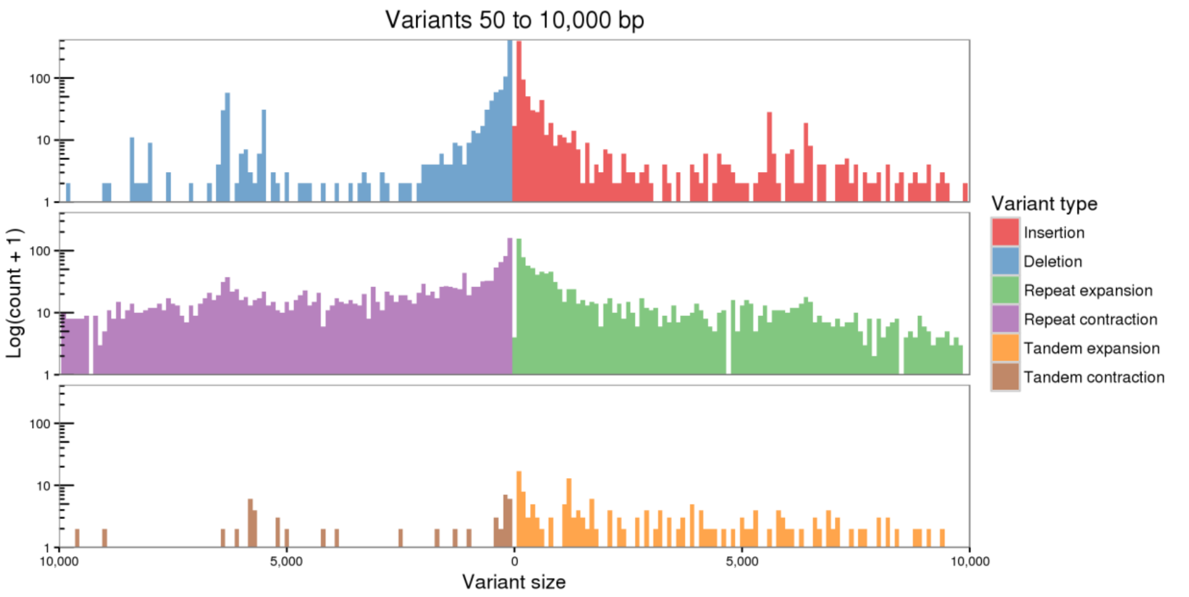


**Figure 3** Structural variation between the two *F. hygrometrica* accessions (Uconn and Zurich) estimated by Assemblytics (http://assemblytics.com/). We used the longer Uconn accession`s assembly as the reference.

#### 4. Whole genome alignment of the *P. patens* and *F. hygrometrica* genomes

We performed whole genome alignment to investigate the genomic divergence between the *F. hygrometrica* and *P. patens* genomes. More specifically, we used minimap2 (Li 2021) with default options to align both *F. hygrometrica* (Zurich and Uconn accessions) and *P. patens* genomes. We then used the generated paf file to extract alignment information employing the R package pafr (https://CRAN.R-project.org/package=pafr). After keeping only primary alignments, we found that 64.10% of the *F. hygrometrica* genome could be covered by homologous *P. patens* sequences. Portions of the *F. hygrometrica* genome with no match from the *P.*

*patens* genomic sequence may represent *F. hygrometrica*-specific genome segments or regions that are too divergent to be alignable. By contrast, *F. hygrometrica* genomic sequences could be only aligned to 37.90% of the *P. patens* genome. Alignable genomic segments showed a mean divergence of 19.08% (IQR=0.56%-42.29%; median= 18.65). Furthermore, only 11.30 % of the alignable regions were nucleotide matches and the rest was covered by mismatches and gaps. This implies that both genomes have obtained highly divergent or genome-specific sequence content and that the absolute and proportional length of genome-specific sequences is about twice as high in *P. patens* than in *F. hygrometrica*.

**A**

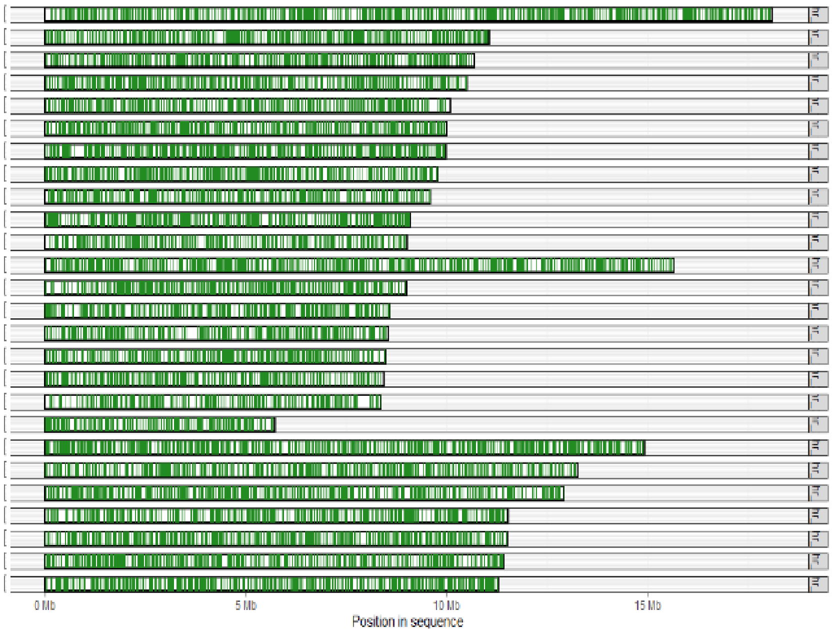

**B**

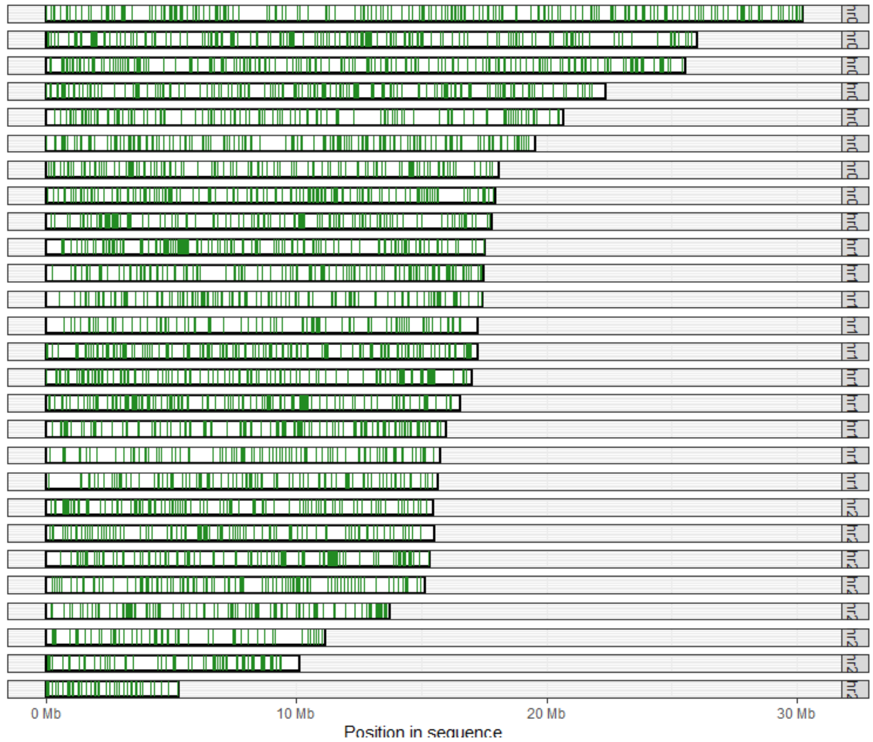


**Figure 4** Visualization of genome mapping between the *F. hygrometrica* Zurich and *Physcomitrium* (*Physcomitrella*) *patens* genomes. (A) *P. patens* genome mapped to the pseudomolecules of the *F. hygrometrica* genome. (B) *F. hygrometrica* genome mapped to the *P. patens* genome. The 26 and 27 pseudomolecules of *F. hygrometrica* and *P. patens* are represented as black framed boxes. Segments of the pseudomolecules covered by valid mappings of the alternate genome`s sequence are shown in green.

#### 5. Gene set comparison of the two *F. hygrometrica* accessions (Zurich and Uconn)

It was reported that comparative annotation of closely related genomes can lead to more accurate gene models. Therefore, a comparative annotation with Augustus in CGP mode was carried out in addition to the Braker2 annotation obtained separately for the Zurich and Uconn accession of *F. hygrometrica*. The softmasked genomes were first aligned using Cactus. The resulting alignment file in hal format was split into chunks with a length of 1 Mb with an overlap of 25 kb, converted to maf format and loaded into a SQlite database. Intron hints were computed separately for each accession using mapped RNAseq data with the bam2hints program included in the Augustus release. Exonpart hints were created for the Zurich accession using RNAseq coverage data in wig format together with the wig2hints utility from Augustus. Both, intron and exonpart hints, were added to the SQlite database as well. The database including the aligned genomes and hints from RNAseq data was then used in a comparative gene annotation run with Augustus in CGP mode. The resulting comparative annotations were then merged with the corresponding BRAKER annotation for each accession using the joingenes program included in the Augustus package. Nevertheless, this approach has led to significantly worse BUSCO scores than that of the individual annotations by fragmenting a significant proportion of high-quality and valid gene models. Therefore, we decided not to use the comparative annotation approach.

Therefore, we created two independent annotations for the Zurich and Uconn accessions of *F. hygrometrica* as described in the methods section of the main text. After visual inspection of the two genome annotations, we found that some gene models predicted on one accession`s genome were apparently missing in the alternative accession`s genome. This could not be alleviated even after several attempts of fine-tuning the parameter space used. Therefore, we decided to consolidate the two accessions` genome annotation manually. To do so, we first identified accession-specific gene models by searching coding sequences (CDS) of the Zurich against the Uconn accession and vice versa using blastn with an e-value threshold of $10^{-6}$ and a minimum query coverage of 70%. This analysis identified 5,211/3,410 Zurich- and Uconn-specific gene models, respectively. After identifying putative accession-specific models, we used their CDS to search for the other accessions genome sequence using exonerate v2.2 (Slater and Birney 2005) using the est2genome option and a minimum coverage and sequence similarity threshold of 70%. Using this approach, only 565 gene models of the Zurich accession`s predicted gene set had no valid hit against the Uconn accession`s genome sequence. Similarly, only 1360 predicted gene models of the Uconn accession had no valid match on the Zurich accession`s genome sequence. These results suggested that many of the accession-specific gene models are false positives due to the stochastic nature of the gene prediction algorithm.

To consolidate the predicted gene sets of the two accessions, we intended to identify and predict the homologous gene models of the putatively accession-specific gene models in the alternate accession`s genome using GenblastA/G v1.39 (She et al. 2009, 2011) (-e 1e-5 -g T -d 10000 -r 5 -c 0.8 -i 15 -x 3 -re 1 -rm 6 -rl 500). The algorithm implemented in GenblastA/G can identify homologous gene structures in the genomic sequence of the alternate strain even in the presence of structural differences (intron/exon number/length). It can also search by extending gene models to find potential start and stop codons. With this approach the number of Zurich- and Uconn-specific gene models decreased dramatically to 52 and 474, respectively. To consolidate both genome annotation, we appended Genblast gene models showing no overlap with gene models of the original annotation to each accession`s genome annotation using bedtools intersect function (Quinlan and Hall 2010). Finally, the appended gene models were filtered using gFACs applying the very same thresholds described in the main text. Therefore, our analysis shows that the actual number of accession-specific genes is relatively low and is comparable to observations made in vascular plants.

#### 6. Phylogenetic analysis of LTR Copia and Gypsy super-families

For further sub-classification of annotated LTR elements of the Copia and Gypsy super-family, we retrieved alignments of reverse transcriptase (RT) domain of several known sub-families from the *Gypsy* Database (Llorens et al. 2011) and built a Hidden Markov Model (HMM) for the protein domain employing the hmmbuild function of the HMMER software package v3.3 (Potter et al. 2018). We then translated nucleotide sequences of Copia and Gypsy elements annotated in the *F. hygrometrica* genome to their respective peptide sequences in all six frames and scanned them for the presence of an RT domain in combination with the previously built HMM using the hmmsearch utility (Eddy 2011) of the HMMER software package v3.3 (Potter et al. 2018). We retained significant hits (E-value threshold: 1e-5) covering at least 80% of the protein domain. We discarded all LTR elements which had multiple valid hits in different reading frames. The remaining RT domains were aligned with MUSCLE v3.8.31 (Edgar 2004) using the consensus sequence of the RT domain of the Bel-Pao superfamily, retrieved from GypsyDB, as outgroup. An optimal tree was inferred from bootstrap tests with 1000 replicates (Felsenstein 1985) using the Neighbor-Joining method (Saitou and Nei 1987) in MEGA X v10.2.5 (Kumar et al. 2018). The phylogenetic tree,classification of TE-elements, and their distribution along the *F. hygrometrica* pseudomolecules are shown on Figures 5-7.

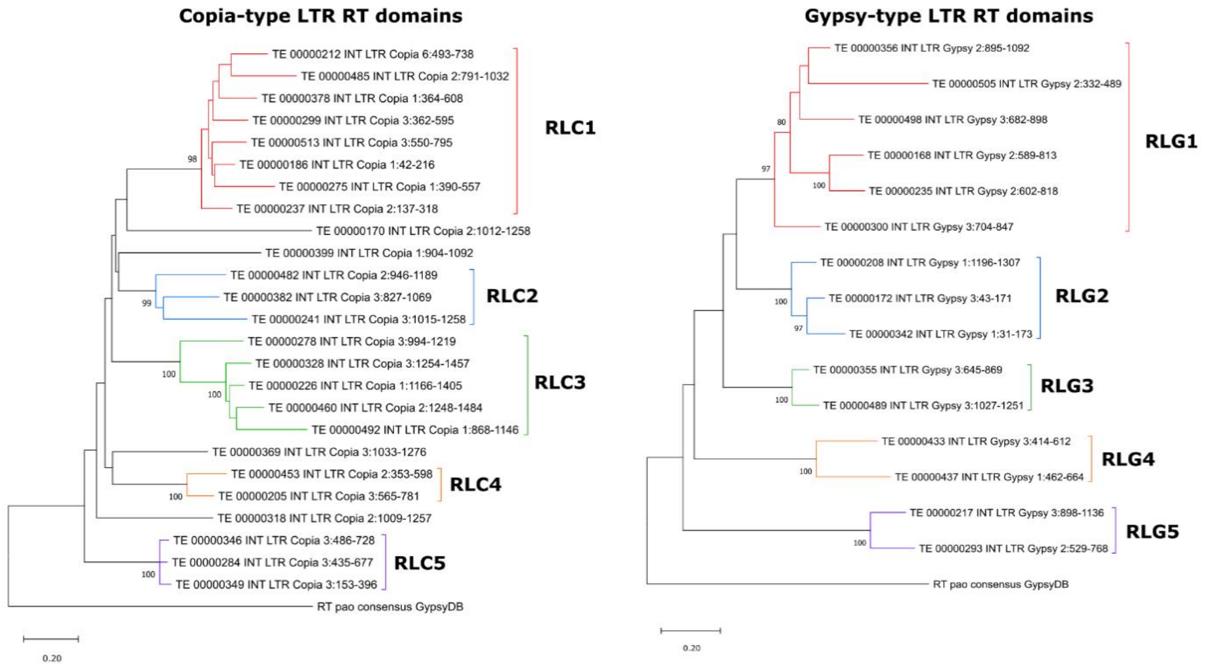


**Figure 5** Phylogenetic trees generated from peptide alignments of the reverse transcriptase (RT) domain of LTR elements from the Copia (left tree) and Gypsy (right tree) superfamilies residing in the *F. hygrometrica* acc. Zurich genome. Bootstrap consensus trees with 1000 replicates (Felsenstein 1985) using the Neighbor-Joining method (Saitou and Nei 1987) are shown. The percentage of replicate trees in which the associated sequences clustered together are shown for highly supported branches. The trees are drawn to scale and represent the evolutionary distances in number of amino acid substitutions per site, which were computed using the Poisson correction method (Zuckerkandl and Pauling 1965). The consensus sequence of the RT domain found in LTR elements of the Bel-Pao superfamily, retrieved from the Gypsy Database release 2.0 (Llorens et al. 2011), was used as an outgroup for the analysis. A temporary family name (RLC1-5 and RLG1-5) was assigned to sequences which clustered repeatedly together. Tree generation and phylogenetic analyses were conducted using MEGA X (Kumar et al. 2018).

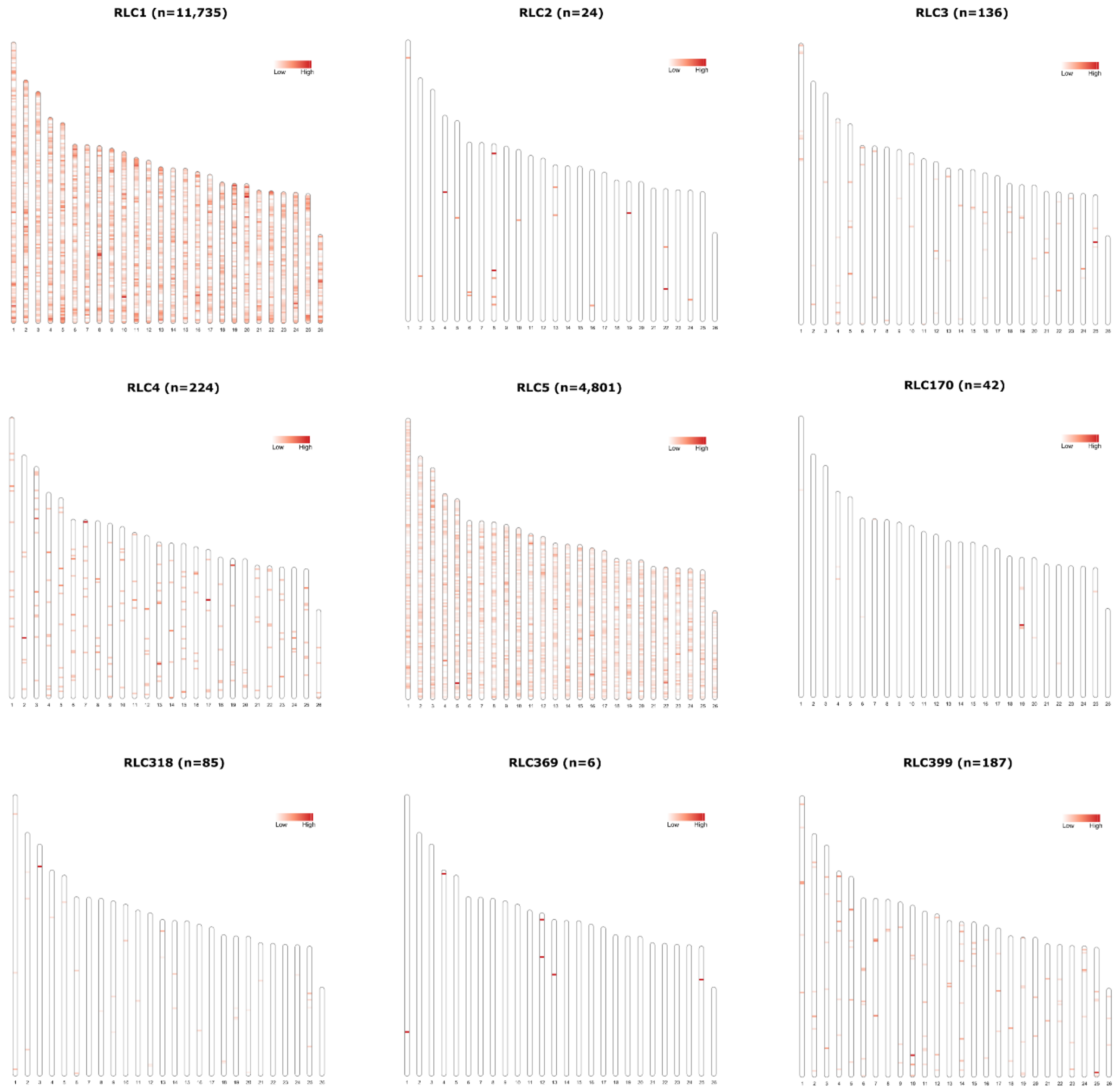


**Figure 6** Distribution of LTR retrotransposons of the Copia superfamily in 100kb windows along chromosome-scale pseudomolecules of the *F. hygrometrica* acc. Zurich genome assembly. Elements with high sequence similarity of the amino acid sequence of their RT domain were grouped together in sub-families (RLC1-5) based on phylogenetic analyses as shown in Figure 1. Elements which could not be assigned to a sub-family are shown individually. Figures were generated with the R package RIdeogram (Hao et al. 2020).

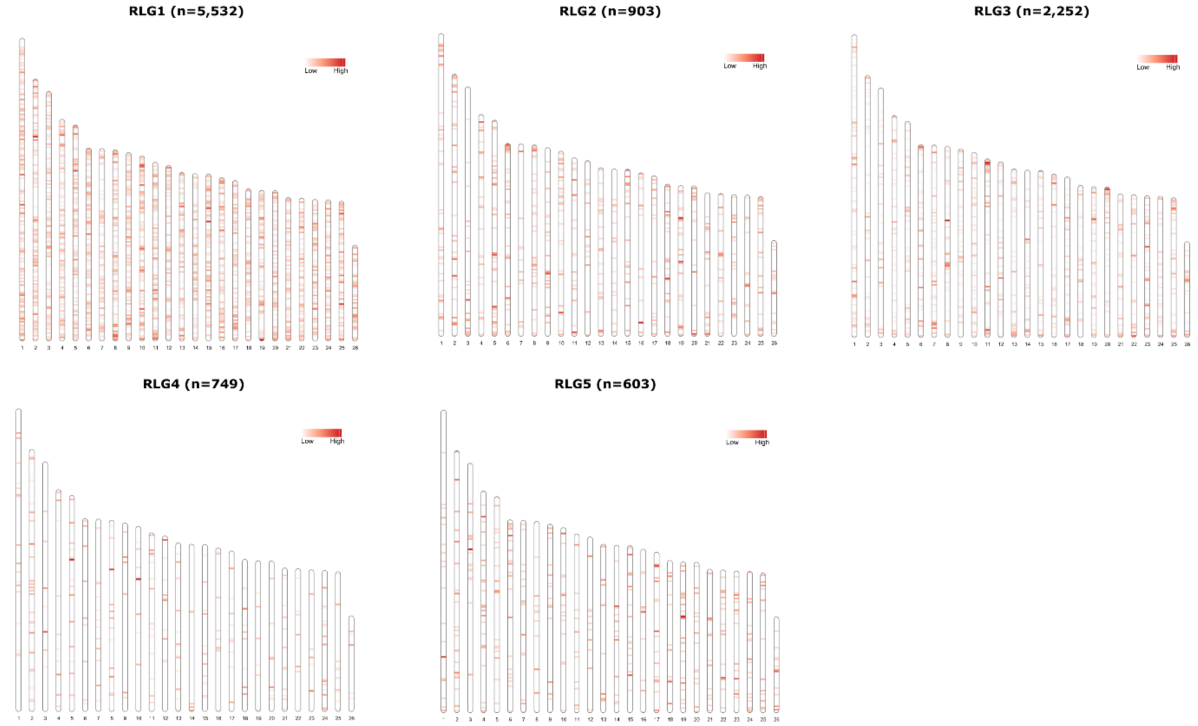


**Figure 7** Distribution of LTR retrotransposons of the Gypsy superfamily in 100kb windows along chromosome-scale pseudomolecules of the *F. hygrometrica* acc. Zurich genome assembly. Elements with high sequence similarity of the amino acid sequence of their RT domain were grouped together in sub-families (RLG1-5) based on phylogenetic analyses as shown in Figure 1. Figures were generated with the R package RIdeogram (Hao et al. 2020).

#### 7. References

Armstrong, Joel, Glenn Hickey, Mark Diekhans, Ian T. Fiddes, Adam M. Novak, Alden Deran, Qi Fang, et al. 2020. “Progressive Cactus Is a Multiple-Genome Aligner for the Thousand-Genome Era.” *Nature* 587 (7833): 246–51. https://doi.org/10.1038/s41586-020-2871-y.

Cove, D. J., P.-F. Perroud, A. J. Charron, S. F. McDaniel, A. Khandelwal, and R. S. Quatrano. 2009. “Culturing the Moss Physcomitrella Patens.” *Cold Spring Harbor Protocols* 2009 (2): pdb.prot5136-pdb.prot5136. https://doi.org/10.1101/pdb.prot5136.

Dolezel, J, J Bartos, H Voglmayr, and J Greilhuber. 2003. “Nuclear DNA Content and Genome Size of Trout and Human.” *Cytometry A.* 51 (2): 127–28. https://doi.org/10.1002/cyto.a.10013.

Durand, Neva C., Muhammad S. Shamim, Ido Machol, Suhas S.P. Rao, Miriam H. Huntley, Eric S. Lander, and Erez Lieberman Aiden. 2016. “Juicer Provides a One-Click System for Analyzing Loop-Resolution Hi-C Experiments.” *Cell*

*Systems* 3 (1): 95–98. https://doi.org/10.1016/j.cels.2016.07.002.

Galbraith, David W., Kristi R. Harkins, Joyce M. Maddox, Nicola M. Ayres, Dharam P. Sharma, and Ebrahim Firoozabady. 1983. "Rapid Flow Cytometric Analysis of the Cell Cycle in Intact Plant Tissues." *Science (New York, N.Y.)* 220 (4601): 1049–51. https://doi.org/10.1126/science.220.4601.1049.

Greilhuber, J., and I Ebert. 1994. "Genome Size Variation in Pisum Sativum." *Genome* 37 (4): 646–55. https://doi.org/10.1139/g94-092.

Greilhuber, J, and E.M. Temsch. 2001. "Feulgen Densitometry : Some Observations Relevant to Best Practice in Quantitative Nuclear DNA Content Determination." *Acta Bot. Croat.* 60 (2): 285–98.

Hickey, Glenn, Benedict Paten, Dent Earl, Daniel Zerbino, and David Haussler. 2013. "HAL: A Hierarchical Format for Storing and Analyzing Multiple Genome Alignments." *Bioinformatics (Oxford, England)* 29 (10): 1341–42. https://doi.org/10.1093/bioinformatics/btt128.

Lang, Daniel, Kristian K. Ullrich, Florent Murat, Jörg Fuchs, Jerry Jenkins, Fabian B. Haas, Mathieu Piednoel, et al. 2018. "The Physcomitrella Patens Chromosome-Scale Assembly Reveals Moss Genome Structure and Evolution." *Plant Journal* 93 (3): 515–33. https://doi.org/10.1111/tpj.13801.

Li, Heng. 2021. "New Strategies to Improve Minimap2 Alignment Accuracy." *Bioinformatics (Oxford, England)* 37 (23): 4572–74. https://doi.org/10.1093/bioinformatics/btab705.

Marçais, Guillaume, and Carl Kingsford. 2011. "A Fast, Lock-Free Approach for Efficient Parallel Counting of Occurrences of k-Mers." *Bioinformatics (Oxford, England)* 27 (6): 764–70. https://doi.org/10.1093/bioinformatics/btr011.

Otto, F. J., H. Oldiges, W. Göhde, and V. K. Jain. 1981. "Flow Cytometric Measurement of Nuclear DNA Content Variations as a Potential in Vivo Mutagenicity Test." *Cytometry* 2 (3): 189–91. https://doi.org/10.1002/cyto.990020311.

Quinlan, Aaron R., and Ira M. Hall. 2010. "BEDTools: A Flexible Suite of Utilities for Comparing Genomic Features." *Bioinformatics* 26 (6): 841–42. https://doi.org/10.1093/bioinformatics/btq033.

Ranallo-Benavidez, T. Rhyker, Kamil S. Jaron, and Michael C. Schatz. 2020. "GenomeScope 2.0 and Smudgeplot for Reference-Free Profiling of Polyploid Genomes." *Nature Communications* 11 (1): 1432. https://doi.org/10.1038/s41467-020-14998-3.

Reski, Ralf, Merle Faust, Xiao-Hui Wang, Michael Wehe, and Wolfgang O. Abel. 1994. "Genome Analysis of the Moss Physcomitrella Patens (Hedw.) B.S.G." *Molecular and General Genetics MGG* 244 (4): 352–59. https://doi.org/10.1007/BF00286686.

She, Rong, Jeffrey S.C. Chu, Ke Wang, Jian Pei, and Nansheng Chen. 2009. "GenBlastA: Enabling BLAST to Identify Homologous Gene Sequences." *Genome Research* 19 (1): 143–49. https://doi.org/10.1101/gr.082081.108.

She, Rong, Jeffrey Shih-Chieh Chu, Bora Uyar, Jun Wang, Ke Wang, and Nansheng

Chen. 2011. “GenBlastG: Using BLAST Searches to Build Homologous Gene Models.” *Bioinformatics* 27 (15): 2141–43. https://doi.org/10.1093/bioinformatics/btr342.

Slater, Guy St C, and Ewan Birney. 2005. “Automated Generation of Heuristics for Biological Sequence Comparison.” *BMC Bioinformatics* 6 (February): 31. https://doi.org/10.1186/1471-2105-6-31.

Temsch, Eva, Johann Greilhuber, and Robert Krisai. 2010. “Genome Size in Liverworts.” *Preslia* 82 (1): 63–80.

Yu, Jun, Songnian Hu, Jun Wang, Gane Ka-Shu Wong, Songgang Li, Bin Liu, Yajun Deng, et al. 2002. “A Draft Sequence of the Rice Genome (Oryza Sativa L. Ssp. Indica).” *Science (New York, N.Y.)* 296 (5565): 79–92. https://doi.org/10.1126/science.1068037.

Felsenstein, Joseph. 1985. “CONFIDENCE LIMITS ON PHYLOGENIES: AN APPROACH USING THE BOOTSTRAP.” *Evolution* 39 (4): 783–91. https://doi.org/10.1111/j.1558-5646.1985.tb00420.x.

Hao, Zhaodong, Dekang Lv, Ying Ge, Jisen Shi, Dolf Weijers, Guangchuang Yu, and Jinhui Chen. 2020. “RIdeogram: Drawing SVG Graphics to Visualize and Map Genome-Wide Data on the Idiograms.” *PeerJ Computer Science* 6 (January): 1–11. https://doi.org/10.7717/peerj-cs.251.

Kumar, Sudhir, Glen Stecher, Michael Li, Christina Knyaz, and Koichiro Tamura. 2018. “MEGA X: Molecular Evolutionary Genetics Analysis across Computing Platforms.” Edited by Fabia Ursula Battistuzzi. *Molecular Biology and Evolution* 35 (6): 1547–49. https://doi.org/10.1093/molbev/msy096.

Llorens, Carlos, Ricardo Futami, Laura Covelli, Laura Domínguez-Escribá, Jose M. Viu, Daniel Tamarit, Jose Aguilar-Rodríguez, et al. 2011. “The Gypsy Database (GyDB) of Mobile Genetic Elements: Release 2.0.” *Nucleic Acids Research* 39 (SUPPL. 1): D70–74. https://doi.org/10.1093/nar/gkq1061.

Saitou, N., and M. Nei. 1987. “The Neighbor-Joining Method: A New Method for Reconstructing Phylogenetic Trees.” *Molecular Biology and Evolution* 4 (4): 406–25. https://doi.org/10.1093/oxfordjournals.molbev.a040454.

Zuckerkandl, Emile, and Linus Pauling. 1965. “Evolutionary Divergence and Convergence in Proteins.” In *Evolving Genes and Proteins*, 97–166. Elsevier. https://doi.org/10.1016/b978-1-4832-2734-4.50017-6.
